# Inferring system-level brain communication through multi-scale neural activity

**DOI:** 10.1101/2020.11.30.404244

**Authors:** Yin-Jui Chang, Yuan-I Chen, Hsin-Chih Yeh, Jose M. Carmena, Samantha R. Santacruz

## Abstract

Fundamental principles underlying computation in multi-scale brain networks illustrate how multiple brain areas and their coordinated activity give rise to complex cognitive functions. Whereas the population brain activity has been studied in the micro-to meso-scale in building the connections between the dynamical patterns and the behaviors, such studies were often done at a single length scale and lacked an explanatory theory that identifies the neuronal origin across multiple scales. Here we introduce the NeuroBondGraph Network, a dynamical system incorporating both biological-inspired components and deep learning techniques to capture cross-scale dynamics that can infer and map the neural data from multiple scales. We demonstrated our model is not only 3.5 times more accurate than the popular sphere head model but also able to predict synchronous neural activity and extract correlated low-dimensional latent dynamics. We also showed that we can extend our methods to robustly predict held-out data across several weeks. The extracted effective connectivity agreed with the neuroanatomical hierarchy of motor control. Accordingly, the NeuroBondGraph Network opens the door to revealing comprehensive understanding of the brain computation, where network mechanisms of multi-scale communications are critical.

## Introduction

Employing system-level dynamics, billions of individual neurons coordinate activities to drive behaviors such as motor preparation^1,2^, motor adaptation^3^, motor timing^4,5^, decision-making^6^, and working memory^7,8^. Current techniques typically make simplified assumptions in which dynamics are linear^9^ or log-linear^10^ to capture neural dynamics by extracting low-dimensional latent processes, and work to date has largely focused on measurements from a single modality. While within-level, nonlinearly correlated neural dynamics encode rich information giving rise to behavior^11^, cross-level nonlinear communication, which uncovers a deeper understanding of system-level neural mechanisms^12,13^, is seldom explored. Since the brain exhibits computational structure across a variety of scales: from single neurons (micro-scale) to functional areas (meso-scale) to cortical networks (macro-scale), a tool that can uncover multi-scale dynamics is critically important for illuminating the mechanistic understanding of brain activity^14^.

Recent cross-level analyses include source localization and cross-level coupling (CLC)^15^. Source localization (e.g., sphere head model^16^) is to identify the brain areas or individual neurons generating the recorded electrical potentials^17^. However, the requirement of high-density recordings, unrealistic assumptions, and uncertainty on conductivity value^18^ limit the fidelity on experimental data. Although CLC has shown the evidence of interactions, no information about how the activity communicates across levels was provided. Here we introduce a multi-scale dynamical model, termed the NeuroBondGraph Network (NBGNet)^19^, to capture the causal interactions between neurons or networks at disparate scales. The NBGNet is constructed based on *a priori* knowledge obtained from a Bond Graph (BG)^20^-derive dynamic systems, enabling us to incorporating biologically realistic assumptions in our model. The recurrent neural network (RNN) framework is then implemented to approximate the nonlinear mapping for characterizing the implicit relations of multi-scale brain activity. Unlike source localization, we bypass the issue of unrealistic assumptions which lead to inaccurate predictions. Compared to CLC, the NBGNet models the causal contributions of cross-scale network. While purely data-driven methods such as black-box RNN may achieve similar performance, the NBGNet approach provides powerful interpretability to evaluate both within- and cross-scale causal interactions.

The NBGNet model is universal in the sense that it can be used for any combination of neural activity at different scales (or even the same scale) with the appropriate selection of the BG. In this work, we use two specific types of simultaneously recorded real data for the purpose of characterizing our method and demonstrating the power of this model. Namely, here we use local field potentials (LFPs) and signals recorded from screw type electrodes implanted in the skull (screw electrocorticography or screw ECoG^21,22^), acquired from a rhesus macaque performing a center-out joystick task. Screw ECoG, rather than electroencephalography, is chosen due to its improved signal-to-noise ratio and stability. The structure of the NBGNet for these two particular data types easily be extended to other field potential signals, and with some modifications can even be utilized for spiking data.

We demonstrate that the presented method performs more accurately in both the forward and inverse problem compared to other techniques. Low error (3.5-fold improvement) and strong similarity (2.2- and 1.6-fold increase) in both time- and phase-domain to ground-truth signals validate the NBGNet as an accurate solution. Furthermore, the NBGNet can capture and accurately reconstruct single-trial low-dimensional neural dynamics. Behavioral variables can also be predicted by NBGNet-inferred activity as accurately as using empirical measurements. Finally, we examine the stability of the performance of the model and reveal that the learned dynamical system maintains the predictive power over several weeks. Last but not least, comparing with the neuroanatomical hierarchy of motor control^23^, the NBGNet-derived causal interactions between channels hold great potential to investigating brain mechanisms across multiple scales.

## Results

### Overview of NBGNet

To introduce the NBGNet (**Fig. 1**), we start with a generic dynamical system, where the evolution of latent variables and the output is described by the nonlinear functions of latent states and corresponding input. Nonlinearity is derived from the BG, a modeling framework used to investigate how energy is transferred among different system components^20^ and has been adopted for system identification and fault diagnosis^24^, based on the translation between two recording modalities (**Supplementary Fig. 1**). Multi-layer perceptron units are adopted to approximate the nonlinear functions, generating a sparse RNN, NBGNet, which maximizes the likelihood of the observed brain signals with its internal states.

**Fig. 1.**
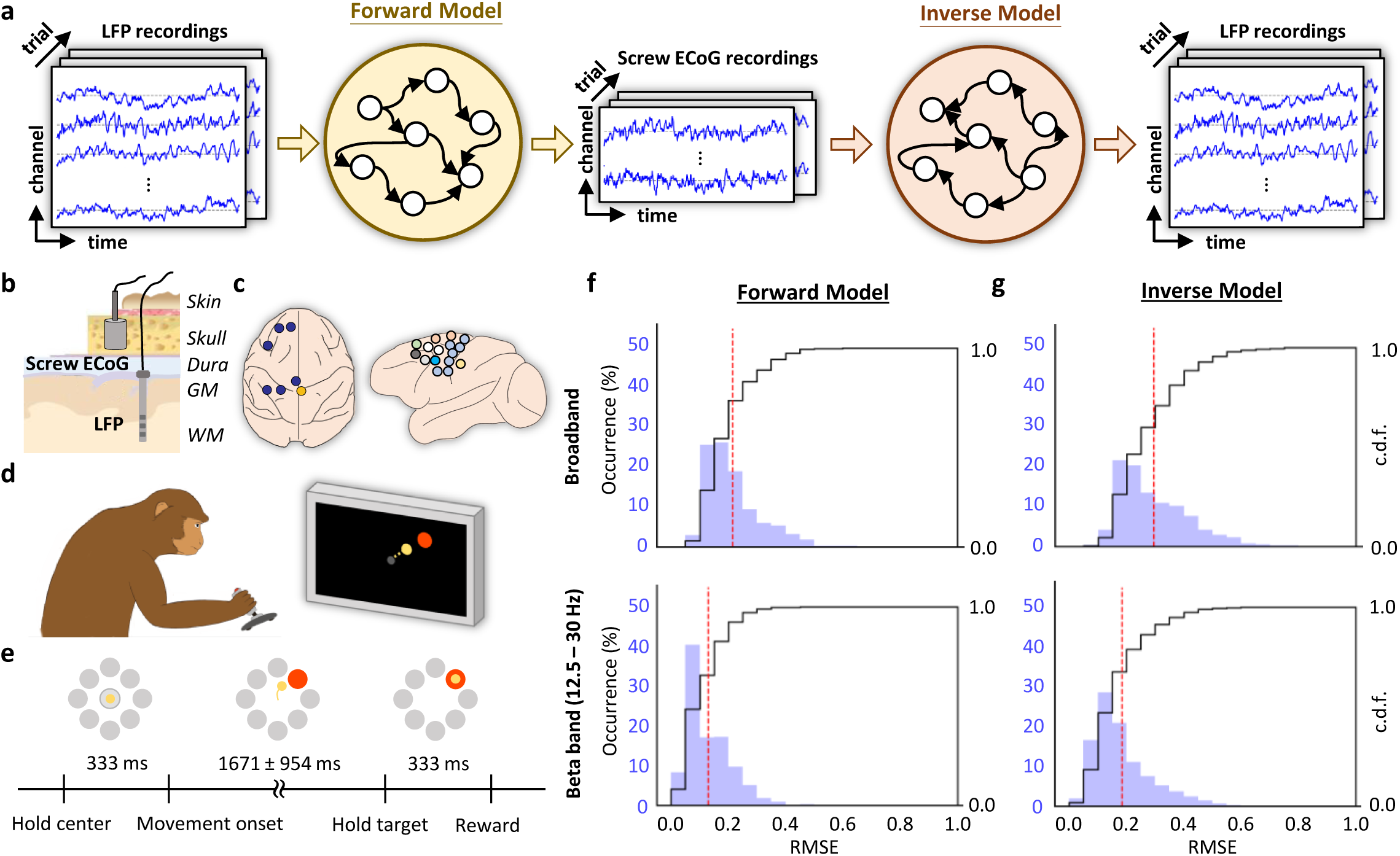
NBGNet is a sparse recurrent neural network that utilizes nonlinear dynamics to model the translation between multimodal brain activities. **a**, Schematic overview of NBGNet architecture for forward and inverse modeling between LFP and screw ECoG. Details are provided in the main text. **b**, Spatial relationships between LFP and screw ECoG. **c**, LFP data was acquired from one hemisphere, and screw ECoG signals were recorded across both hemispheres. **d**, Monkey performed a center-out reaching task using a joystick. **e**, Schematic of protocol for the experiments. **f,g**, Histogram and cumulative distribution function (c.d.f.) of RMSE in broadband and beta band (12.5 – 30 Hz) for forward model (**f**) and inverse model (**g**; red dashed line: mean).

The forward solution, forward-NBGNet, models the single-trial screw ECoG as a nonlinear recursive mapping from the LFP (**Fig. 1a-b**). The network’s units to approximate such a mapping depend on three elements: a trial-specific initial state, input signals, and the parameters defining the connections of the model. To mimic the real-time modeling and abide by causality constraints, the network only runs through the trial backward for estimation. The inverse model, inverse-NBGNet, is then developed by inverting forward-NBGNet. The resulting network predicts LFP across time from the temporal sequence of screw ECoG (**Fig. 1a**). As inverse computation is an ill-posed problem which can lead to non-unique and unstable solution^25^, we expect a relatively poorer performance when compared with the forward model. However, NBGNet still provide accurate and reliable estimations.

We trained the NBGNet on simultaneously recorded LFP data from the left hemisphere and screw ECoG data from both hemispheres. This data was recorded while a monkey performed a center-out joystick task (**Fig. 1c-d**; **Supplementary Table 1-2**). We chose 16 LFP channels and 7 screw ECoG channels from distinct areas to obtain a subset of anatomically spatially distributed signals. The monkey began each trial by holding a cursor at the center target. Then, one of eight outer targets was presented on the screen. After 333 ms, the monkey was instructed to move the cursor to the target for rewards. LFP data was recorded using a semichronic multielectrode array (GrayMatter Research, Bozeman, MT) and screw ECoG data was recorded using screw-type electrodes. The length of each trial is variable, but on average is 1.53 ± 1.22 seconds (mean ± s.d.; **Fig. 1e**).

We next analytically validated NBGNet-inferred neural activities by calculating root mean squared errors (**Fig. 1f-g**); reproducing features seen in common neuroscientific analyses (cross-correlation; **Fig. 2**, phase synchrony; **Fig. 3**); reconstructing low-dimensional latent dynamics (**Fig. 5**); inferring details of behavior (**Fig. 6**); and predicting out-of-sample conditions (**Fig. 7**). For all results in this paper, we trained NBGNets without any information about task conditions or behavioral parameters (e.g., real kinematics or eye-tracker data).

**Fig. 2.**
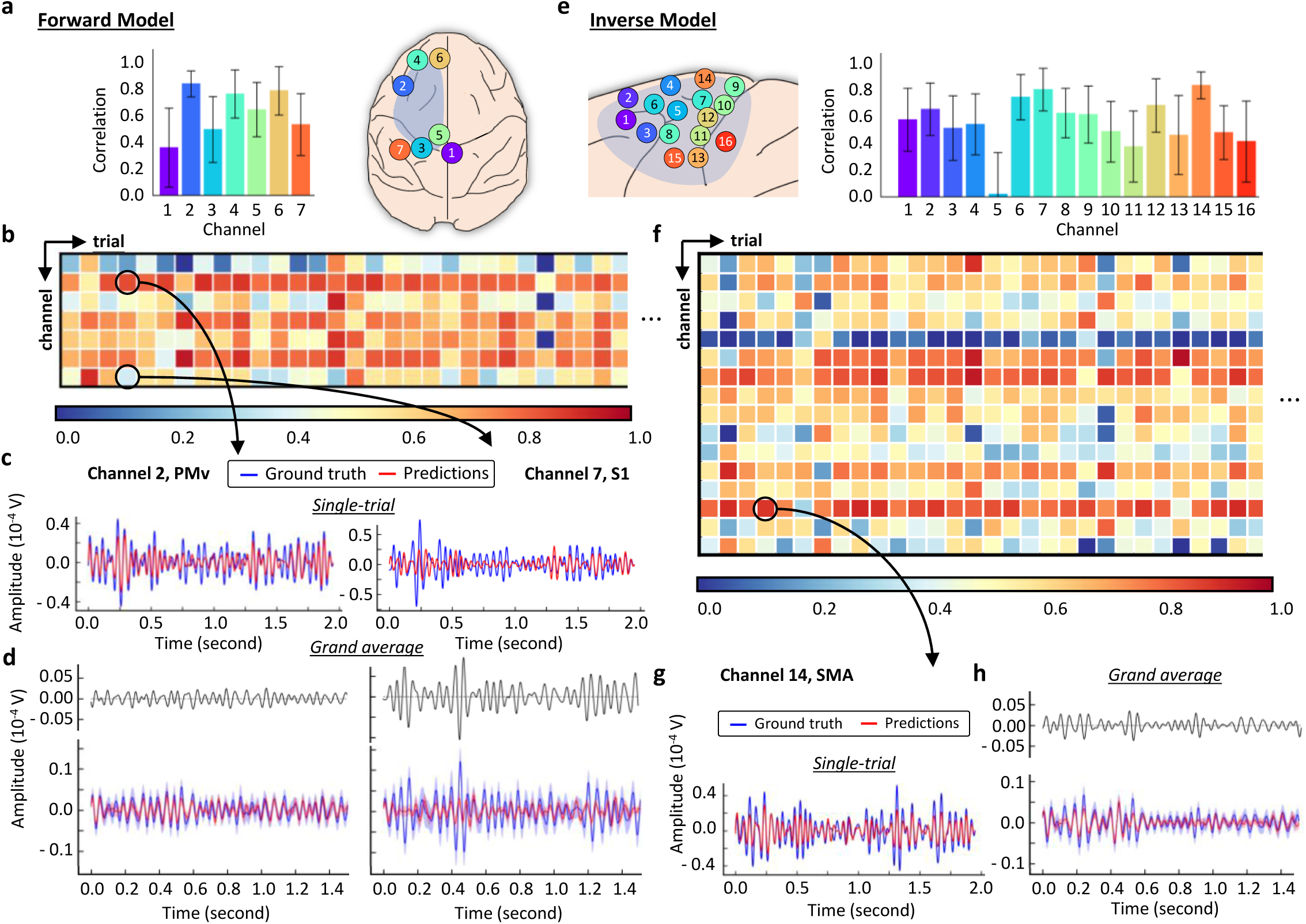
Cross-correlation analysis indicated the similarity between NBGNet inference and ground-truth recordings. **a**, Average correlation coefficient across all the trials (error bars, s.d.; n = 150). Screw ECoG electrodes layout labeled with the channel number. Blue shaded area represents the coverage of LFP channels. **b**, Heatmap showing correlation coefficient for each trial (*x* axis) and each targeted screw ECoG channel (*y* axis). The color key from blue to yellow to red indicates the coefficient from low (0.0) to high (1.0). **c,d**, Two screw ECoG channels (2: PMv, 7: S1) were selected for single trial-based comparison (**c**): ground truth (blue trace) versus model prediction (red trace) in the 4^th^ trial, and grand average-based comparison (**d**): ground truth (blue trace; mean ± s.e.m.) versus network output (red trace; mean ± s.e.m.) and the corresponding error trace (black trace). **e**, Same as **a** for the inverse model, where LFP electrodes are within the coverage of the blue shaded area. **f**, Same as **b** for the inverse model, where *y* axis represents the targeted LFP channels. **g,h**, Same as **c,d** for representative comparison for the inverse model, where only one LFP channel (14: SMA) was chosen.

**Fig. 3.**
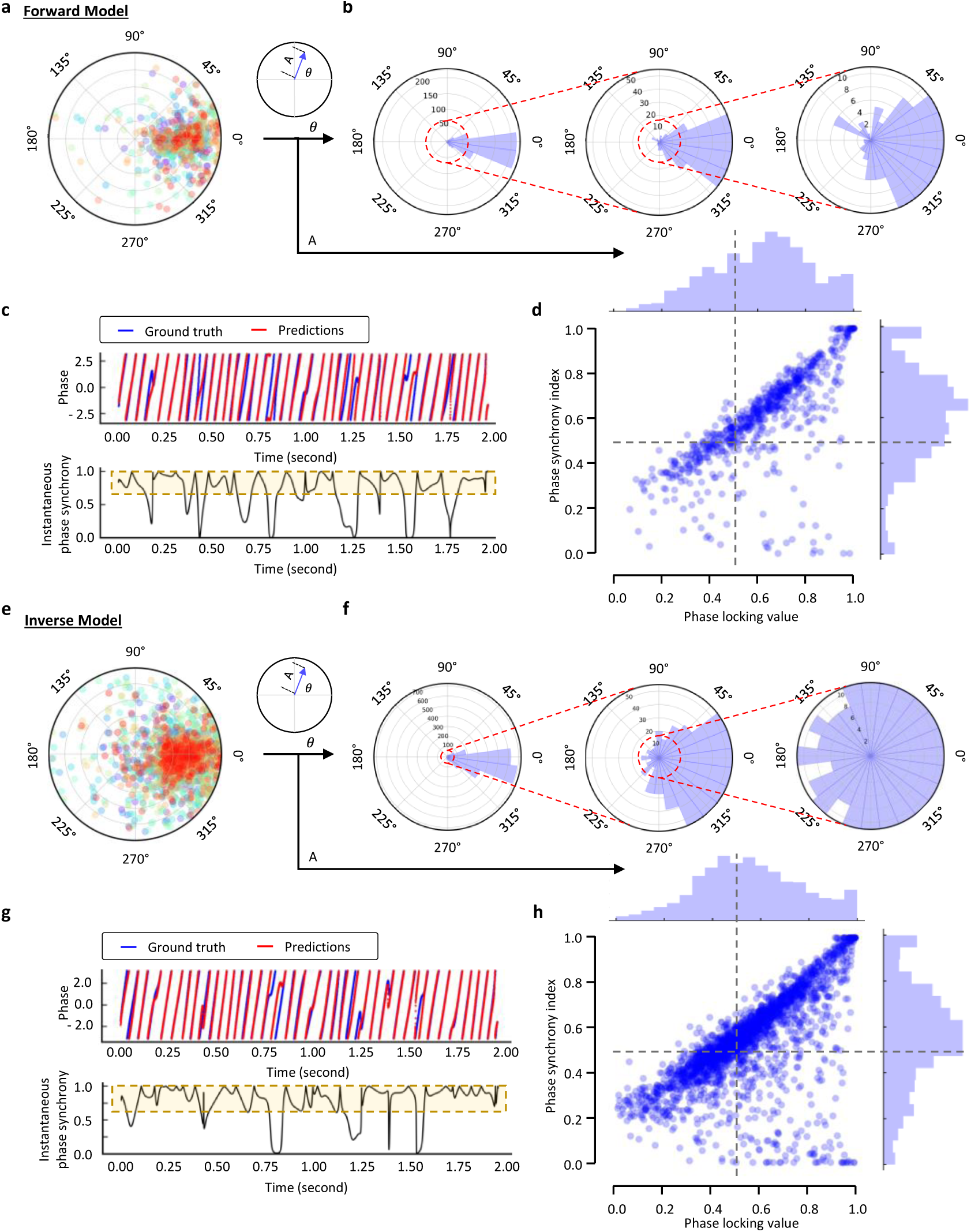
Strong phase synchrony between NBGNet estimations and the experimental recordings. **a**, Polar plots of the mean phase difference averaging across time in each trial for the forward model (n = 1,050). **b**, Angular and corresponding zoom-in histogram of the phase of phase-locking derived from (**a**). **c**, A screw ECoG channel (2: PMv) was selected for demonstrating that NBGNet made predictions in sync with the ground truth in the 4^th^ trial. Instantaneous phase of the ground truth (blue trace) and the model inference (red trace) at each timepoint (upper) was employed to obtain the instantaneous phase synchrony (lower; black trace) across the time. Yellow area showed a strong synchronization utilized to compute the phase synchrony index (PSI). **d**, A scatter plot of phase analysis on each channel and each trial, respectively (n = 1,050), revealing the expected and hidden relations between PSI and phase-locking value (PLV). Histograms of both PLV and PSI are represented on the *x* and *y* axes, respectively. 0.5 was set as thresholds for both PSI and PLV (black dashed line) to identify strong, medium, and poor synchrony regions (**Supplementary Fig. 6**). **e**, Same as **a** for the inverse model (n = 2,400). **f**, Same as **b** for the inverse model, where the histogram was derived from **e. g**, Same as **c**, where the chosen LFP channel for demonstration was the same as **Fig. 2g,h. h**, Same as **d** for the inverse model (n = 2,400).

**Fig. 4.**
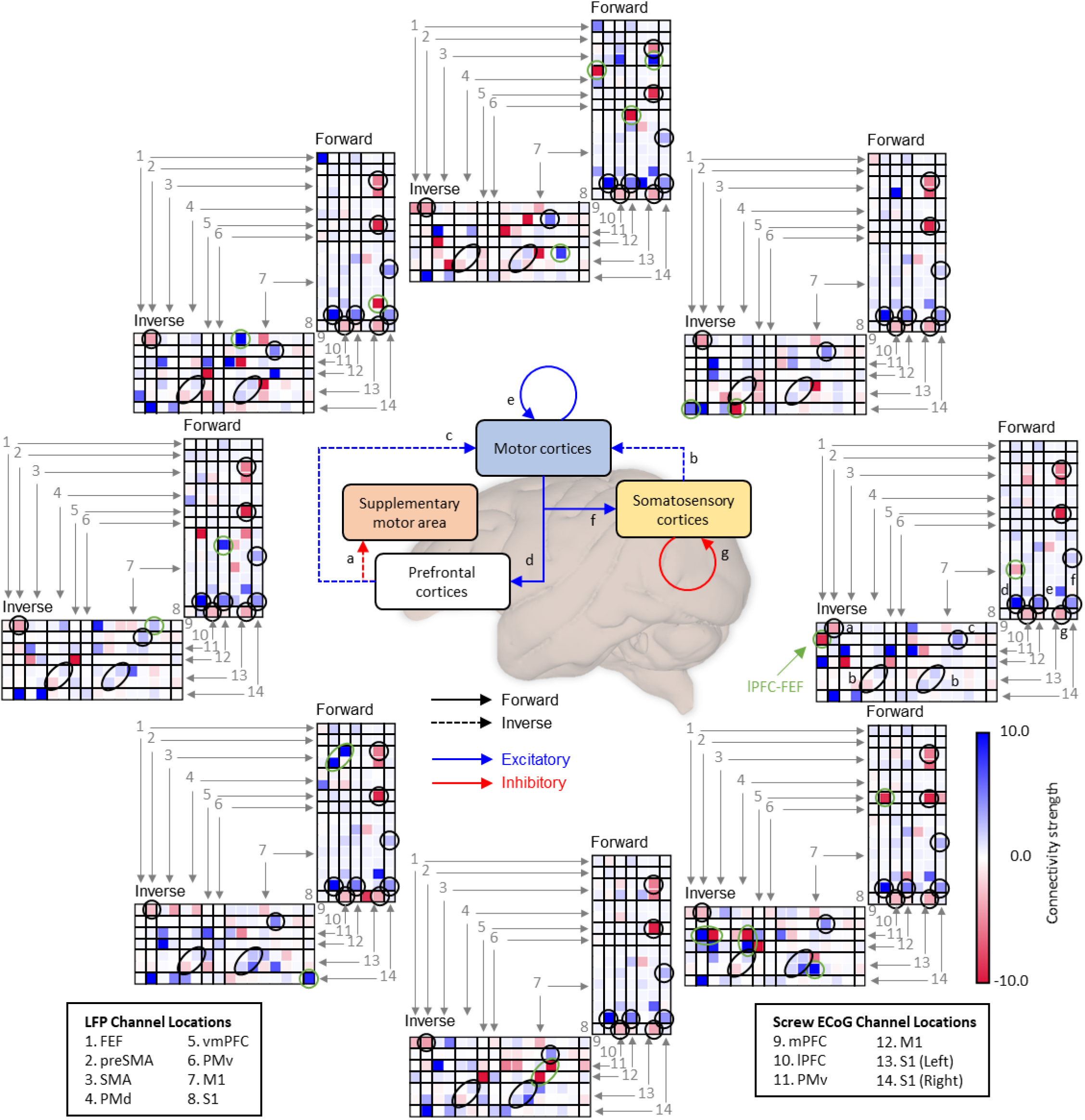
Bi-directional effectivity connectivity extracted from NBGNets exhibited unique and shared patterns in the center-out joystick task. The bi-directional effective connectivity for each target direction was obtained from the NBGNet’s parameters and was averaged over the trials reaching the same target. Each subfigure corresponds to a target position. The vertical axes represent the channels where the connection originates; the horizontal axes represent the channels where the connection contributes to. The shared patterns were indicated with black circles; the unique patterns were indicated with the green circles. The circuitry diagram (middle) depicts the hierarchical interactions between brain regions from the shared patterns of effective connectivity.

**Fig. 5.**
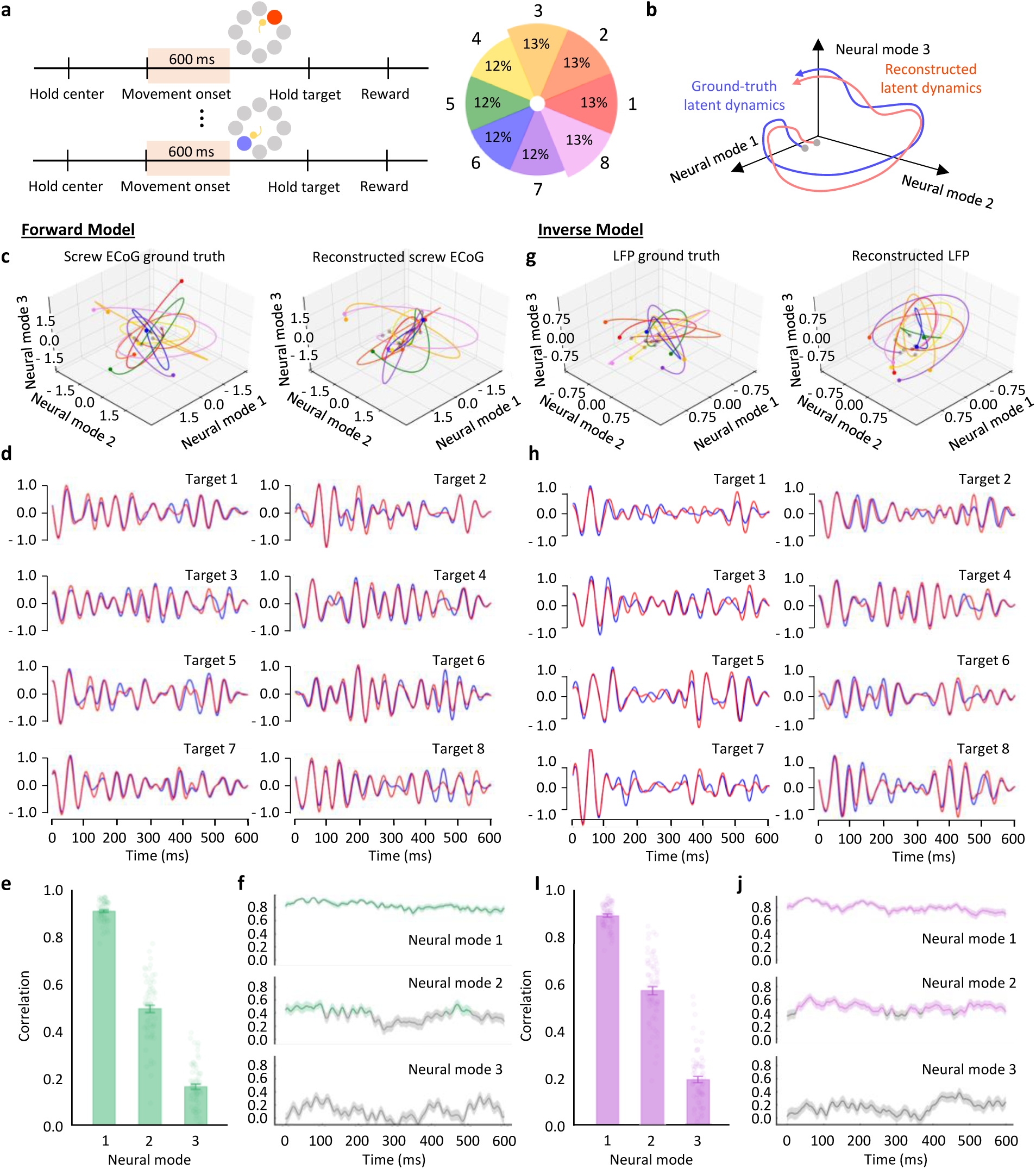
NBGNet captures and reconstructs the latent dynamics in the reaching-out task. **a**, Schematic of protocol indicates the time window used for analysis. Probability of each target direction is uniform. **b**, We predicted that the latent dynamics can be recovered. **c**, Representative latent trajectories derived from the ground-truth screw ECoG (left) and reconstructed screw ECoG (right). Each color represents each target direction in **a. d**, Projection of average ground-truth (blue trace) and reconstructed (red trace) latent trajectories for each target on the first mode. **e**, Bar plot showing the strong magnitude of the correlations between the ground-truth and reconstructed latent trajectories (error bars, s.e.m.; n = 68). **f**, Temporal correlation trajectories for each neural mode (green trace when above the threshold as 0.4; grey trace as below the threshold; mean ± s.e.m.). **g**, Same as **c** for the inverse model to reconstruct the latent trajectories derived from LFPs. **h**, Same as **d** for the projection of average ground-truth LFPs-derived (blue trace) and reconstructed LFPs-derived (red trace) latent trajectories. **i**, Same as **e** for the correlation between the latent trajectories obtained from recorded LFPs and estimated LFPs. **j**, Same as **f** for the inverse model (purple trace when above the threshold as 0.4; grey trace as below the threshold).

**Fig. 6.**
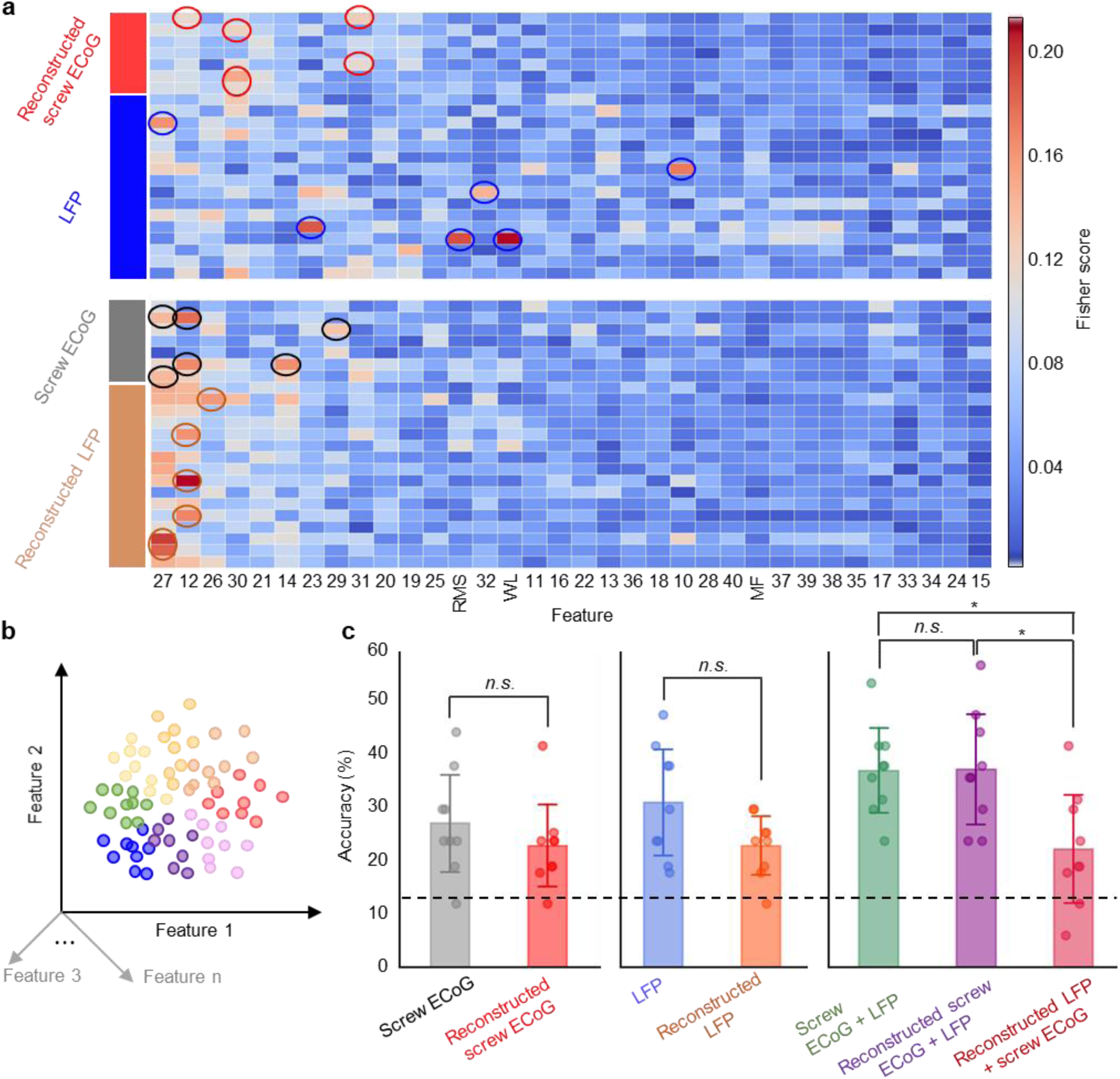
NBGNet inference can be used to predict the movement behavior. **a**, Heatmap showing Fisher score for each feature (*x* axis) and each channel (*y* axis). Chose features for classification were highlighted by circles. **b**, The single-trial features were projected into the feature space, where the LDA classifier found the optimal hyperplanes to separate these data points. **c**, Bar plot showing the classification accuracy for each dataset (dashed line, chance performance; error bars, s.d.; n = 5). *p < 0.05 using two-sided paired T-test. *n*.*s*. indicates no significant difference.

**Fig. 7.**
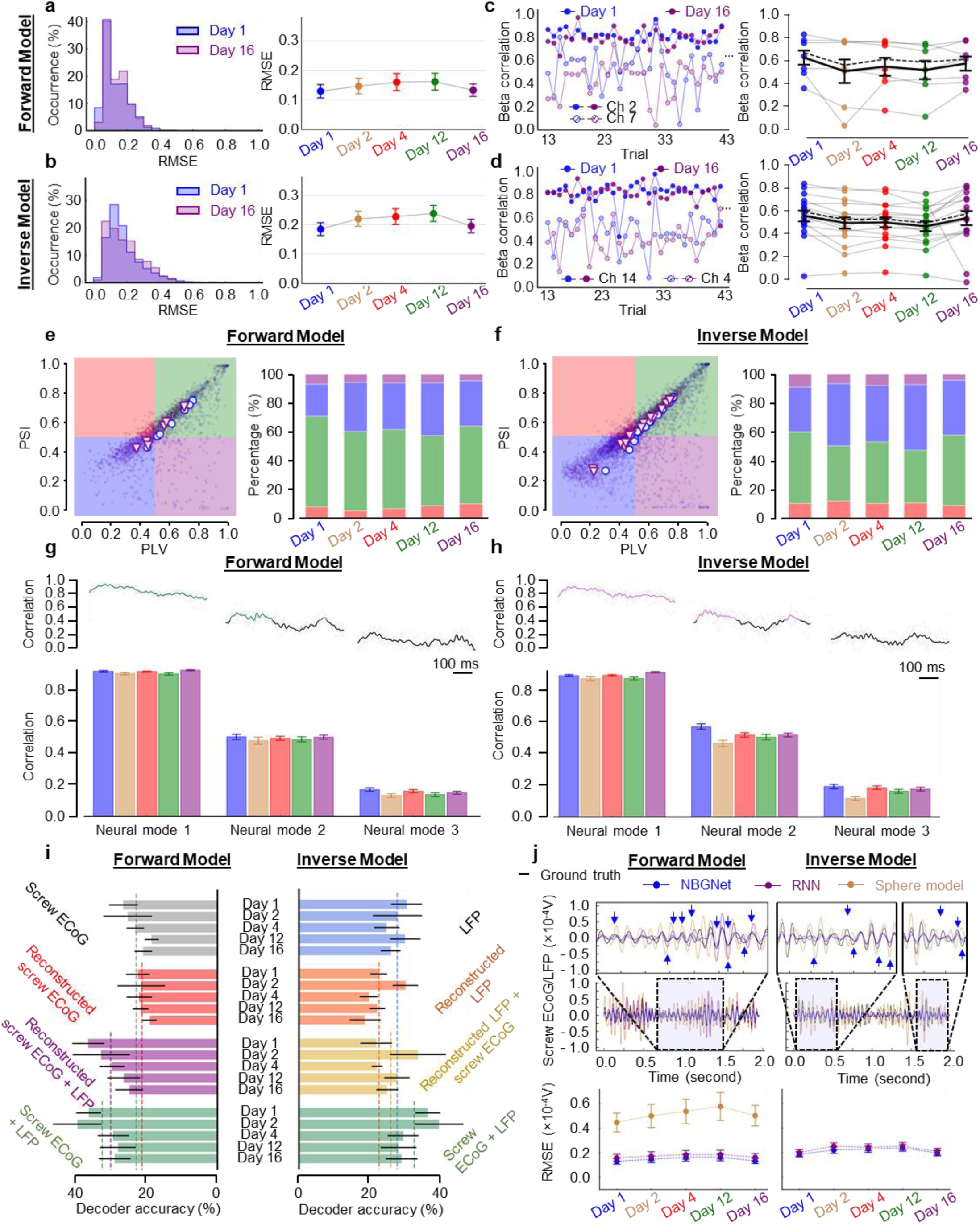
Stability of NBGNet’s predictions for multiple days. **a,b**, Histogram of RMSE (left) at Day 1 (blue) and 16 (purple) for the forward (**a**) and the inverse (**b**) model. Scatter plot of average RMSE (right) showing no significant difference (error bars, s.e.m.; n = 7 and 16 for **a** and **b**). **c,d**, Beta correlation (left) at Day 1 (blue) and 16 (purple) for the forward (**c**) and the inverse (**d**) model. Scatter plot of average beta correlation, where black solid line is obtained by averaging over the channels and black dashed line left the poorest channel out (error bars, s.e.m.; n = 7 and 16 for **c** and **d**). **e,f**, Scatter plot of PSI versus PLI (left) at Day 1 (blue circle) and 16 (purple triangle) for the forward (**e**) and the inverse (**f**) model. Stacked bars (right) demonstrate the percentage of predictions locating in each section. **g,h**, Temporal correlation averaging across days (upper), where colored segments represents stronger correlation as compared with the grey counterparts. Bar plot of average correlation (lower) exhibiting stable performance (error bars; s.e.m.; n = 68, 49, 128, 78, 135 at Day 1, 2, 4, 12, 16). **i**, Bar plots showing the classification accuracy of linear classifier to predict the target direction (error bars, s.d.; n = 5). Dashed line represents the average performance across days. **j**, Performance was quantified by RMSE (bottom) between the ground-truth and inferred signals. Single-trial comparison (top) also demonstrated NBGNet provided the highest prediction accuracy. Blue arrows highlight the time when the NBGNet outperformed other methods. Scatter plot of average RMSE obtained from different methods across days showing no difference in the comparison between each other (error bars, s.e.m.; n = 7 for forward and n = 16 for inverse model). Sphere head model failed to provide predictions within the reasonable range in some trials, thus being excluded for inverse model.

### Validation of NBGNet predictions using a center-out joystick task

The representative dataset used in **Fig. 1** consisted of 150 individual reach trials (average 175 trials/session). Since the beta frequency band (12.5–30 Hz) is strongly implicated in motor behaviors^26,27^, (**Supplementary Fig. 2**), we examined performance with root mean squared error (RMSE) for the predicted broadband signals as well as specifically for the beta band activity extracted from the predictions (**Fig. 1f**). Long short-term memory based recurrent neural network (RNN) was utilized as baseline for quantitative comparison. NBGNet-reconstructed screw ECoG had better match with the ground-truth screw ECoG (RMSE = 0.21 ± 0.09 for broadband; 0.13 ± 0.06 for beta band; mean ± s.d. in the unit of 10^−4^ V) than RNN’s predictions (RMSE = 0.29 ± 0.09 for broadband; 0.16 ± 0.08 for beta band). Additionally, session-averaged RMSE (0.45 for NBGNet < 0.53 for RNN in broadband; 0.15 for NBGNet < 0.17 for RNN in beta band; **Supplementary Fig. 3**) indicated that forward-NBGNet successfully estimated the neural oscillations recorded from screw ECoG.

We also tested whether inverse-NBGNet could recover LFP with screw ECoG recordings (**Fig. 1g**). Similarly, inverse-NBGNet (RMSE = 0.29 ± 0.11 for broadband; 0.18 ± 0.10 for beta band) outperformed RNN (RMSE = 0.35 ± 0.16 for broadband; 0.20 ± 0.09 for beta band). Evidenced with the fact that session-averaged RMSE from NBGNet (0.56 for broadband; 0.21 for beta band) was less than that from RNN (0.59 for broadband; 0.25 for beta band), inverse-NBGNet was able to transform the lower-dimensional screw ECoG into the higher-dimensional LFP.

### NBGNet outputs correlate with ground truth signals

Similarity of oscillation dynamics is another evaluation of integrity of predicted signals using cross-correlations computed on a single-trial single-channel basis. From the representative session (**Fig. 2a-b**), model-predicted signals from all the channels, excluding channel 1 (S1, right hemisphere), were moderately to strongly correlated with the ground-truth signals with average correlation greater than 0.4^28,29^. Strong correlation (correlation coefficient > 0.6) was found on 57% of channels. Interestingly, due to movement-induced activation, channels at anterior brain regions exhibited larger correlation than those at posterior brain regions. In the representative trial, the predicted screw ECoG matched well with the raw screw ECoG for channel 2 (**Fig. 2c**). The grand average also indicated that forward-NBGNet accurately reconstructed screw ECoG with consistently small error trace (**Fig. 2d**). We noted that in channel 7, the performance was relatively poor in the first 0.5 s when the subject was searching for the correct direction of cursor’s movement; however, the performance became better in the rest of time when the direction of movement aligned with the target direction.

We also studied the correlation between the inverse-NBGNet-inferred LFPs and the ground-truth LFPs. Channel 14 (SMA) provided the highest correlation as 0.83 ± 0.10 (mean ± s.d.; **Fig. 2e-f**). Due to unexpectedly larger amplitude, channel 5 (PMd) exhibited no correlation (0.02 ± 0.31). However, the predicted signals on the rest of the channels, excluding channel 11 (M1), were at least moderately correlated with ground-truth activity. 44% of channels showed strong correlations. As more ventral brain region was further from the surface where screw ECoG was recorded, the channels in these regions (channel 11, 13, 15, and 16) showed smaller correlations. In contrast, accurate prediction at channel 14 was expected (**Fig. 2g-h**). Both single-trial- and grand average-based evaluations demonstrated small deviations between the predictions and the ground truths. Taken together, the correlation analysis confirmed the NBGNet’s ability to capture the beta-frequency dynamic features.

### Phase agreement in beta band during movement

Phase-domain coherence is an important tool to determine the integrity of the functional connectivity in brain networks. Accordingly, we examined whether the predicted and the recorded signals were phase-synchronized. Phase-locking value (PLV) has been widely used to measure the inter-trial variability of phase difference, where 1 represents no change in phase difference and 0 reflects the opposite case^30,31^. To assess the intra-trial variability, we adapted PLV by averaging the phase difference across the time rather than the trials (**Fig. 3a,e**). The phase of phase-locking, which indicates the average phase difference, was taken into consideration as well. Instantaneous phase synchrony (IPS) was then used to measure the phase similarity at each time point. Here we utilized a phase synchrony index (PSI) metric which was determined by the fraction of the time when the IPS was greater than a threshold at which the phase difference was less than 45° (threshold = 0.62; **Supplementary Fig. 4a**). If the phase difference seldom exceeds 45°, PSI is close to 1; it is close to zero otherwise. Forward-NBGNet-predicted signals were in sync with the ground truths (73% of average phase difference < 22.5°; average PLV = 0.62; average PSI = 0.60; **Fig. 3b**). A representative channel, channel 2, was demonstrated to generate phase-synchronized predictions (PSI = 0.74; **Fig. 3c**). Since PLV and PSI provided disparate aspects, we assessed the phase synchrony for each channel and each trial in the form of a scatter plot (**Fig. 3d; Supplementary Fig. 4b**). Segmentation of the scatter plot enabled us to study the further details. 78% of predictions exhibited moderate or strong phase synchronization. We next evaluated the inverse-NBGNet’s ability to infer the synchronous LFPs. The predictions were in sync with the ground-truth LFPs (62% of average phase difference < 22.5°; average PLV = 0.56; average PSI = 0.55; **Fig. 3f**). Highly synchronized predictions at a representative channel, channel 14, were observed (PSI = 0.84; **Fig. 3g**). Furthermore, approximately half of the predictions had strong synchronization (**Fig. 3h**). Our phase analysis comprehensively validated that the model predictions are phase-synchronized with the ground truth.

### NBGNet reveals cross-scale causal interactions among brain regions

The complex coordination of brain functions, such as vision, motor preparation, and attention requires the control of causal interactions between areas^32^. Effective connectivity, which represents the influence that a neural system exerts over another^33^, is thus a powerful measure to evaluate the brain computations. We derived the cross-scale effective connectivity from the parameters of the NBGNet, which depicts how the latent states of sources change those of targets. Positive and negative connection strengths correspond to excitatory and inhibitory effects, respectively. The cross-scale effective connectivity for different movement directions in the center-out joystick task exhibited patterns of visual feedback (unique to target position) and voluntary movement (shared across target position; **Fig. 4**). Unique inverse connectivity from lateral prefrontal cortex to frontal eye field was observed during rightward movement, which is consistent with neural activity exhibiting a strong preference for contralateral visual space^34^. Furthermore, as reported in the previous literature^23,35^, we also observed several shared causal interactions (e.g., prefrontal cortex-supplementary motor area, prefrontal cortex-motor cortex, and somatosensory cortex-motor cortex) over all the target directions, aligning well with the abstraction of the hierarchical anatomy of the mammalian nervous system. Thus, NBGNet-derived effective connectivity can illuminate the cross-scale computations underlying brain functions.

### NBGNet captures latent dynamics

Since low-dimensional latent dynamics have been widely used to illuminate the relationship between neural population activity and behavior^36–40^, we also tested whether NBGNet could capture latent dynamics. The window of interest starting from movement onset and ending 600 ms after movement onset was selected, and there is no issue of imbalance target directions (**Fig. 5a**). We computed the neural manifold and the latent dynamics within it using principal component analysis (PCA)^41,42^. The resulting three PCs, termed neural modes, captured the majority of the variance (**Supplementary Fig. 5**), and were used to define the axes of the neural manifold. Canonical correlation analysis (CCA)^43–45^ was performed to align the latent dynamics (**Fig. 5b**). Correlation analysis (Pearson’s *r*) was then utilized to quantify the similarity of these latent dynamics. Since canonical correlations were sorted from largest to smallest, we expected the same trend in the evaluation.

Representative single-trial and grand-average latent trajectories for ground-truth and reconstructed screw ECoG were quite similar for all the target directions (**Fig. 5c,d**). Prediction-derived latent dynamics exhibited strong and moderate correlations with the ground-truth-based counterparts for neural mode 1 (0.91) and mode 2 (0.50), respectively (**Fig. 5e**). Temporal correlation showed the instantaneous performance, allowing us to identify the time-dependency of similarity. We noticed a consistently strong correlation for mode 1 and a strong correlation during the first 250 ms for mode 2, where a drop occurred, followed by a bounce back to correlated level (**Fig, 5f**). Even though, neural mode 2 still exhibited high correlation across the time (41% of the time showed the correlation > 0.4). These observations held for inverse-NBGNet. Latent trajectories derived from inverse-NBGNet-inferred LFPs and the ground-truth LFPs were highly correlated (**Fig. 5g**). Similarly, grand-average latent traces for the first neural mode were almost the same for all the targets (**Fig. 5h**), and a strong and moderate correlation was observed in neural mode 1 (0.89), and 2 (0.57; **Fig. 5i**). Interestingly, the instantaneous correlation was consistently strong over the time interval (100%, and 84% of the time > 0.4 for neural mode 1, and 2; **Fig. 5j**). We noted that the stronger correlation was associated with the higher ratio of variance the neural mode explained. Thus, we demonstrated that the NBGNet could capture the latent dynamics.

### Performance of linear decoder with NBGNet estimations

To understand the information encoded within neural populations, decoding cortical activity is of particular interest^44^. We wondered how accurately linear decoders trained with the model-inferred neural activities would perform. We first extracted candidate features from the dataset and picked six of them using Fisher score^46^ (**Fig. 6a**). Linear discriminant analysis (LDA)^46^ classifiers were then trained with selected features (**Fig. 6b**) to predict the direction of cursor’s movement, where the classification accuracy was evaluated using 9-fold cross-validation.

Candidate features were arranged in descending order based on Fisher score averaging across all the channels (**Fig. 6a**). LDAs were trained with seven conditions: (1) screw ECoG only, (2) reconstructed screw ECoG only, (3) LFP only, (4) reconstructed LFP only, (5) screw ECoG + LFP, (6) reconstructed screw ECoG + LFP, and (7) screw ECoG + reconstructed LFP. Six features were selected for classifiers 1 – 4; while twelve features were selected for classifier 5 – 7. Although the classification accuracy of NBGNet-inferred screw ECoG activity (23%) appears lower than that of ground-truth screw ECoG (27%), there was no significant difference (p = 0.33; **Fig. 6c left**). Similarly, the classification performance between the real and predicted LFPs exhibited no significant difference (p > 0.05; **Fig. 6c middle**), indicating that NBGNet’s inference maintained the discriminant power. As expected, the classifier trained with LFP and screw ECoG outperformed the other conditions (**Fig. 6c right**). Surprisingly, when forward-NBGNet’s predictions were included, the decoding capability reached the same level as using both real signals. A less accurate classification was observed when inverse-NBGNet’s predictions were included as compared with other conditions using both signals (22% for screw ECoG + reconstructed LFP, 37% for both screw ECoG + LFP and reconstructed screw ECoG + LFP), which may result from the less accurate LFP predictions. Together, we demonstrated that the presented model still maintained the integrity of information represented by the neural activity and thus held great potential to improve decoding capability by incorporating appropriate data preprocessing, feature selection, and decoder.

### Stable performance across days

As experiments are often performed across multiple sessions, we tested the stability of NBGNet using the same metrics (**Supplementary Figs. 3, 6-8**). We would like to emphasize that the NBGNet was trained on day 1 and remained fixed for testing in different days. First, the average RMSE between the NBGNet inference and recorded signals for both forward and inverse model were consistent over weeks to a degree almost indistinguishable from that in Day 1 (**Fig. 7a-b**). As was the case for RMSE, with a few individual trial exceptions, the forward- and inverse-model beta correlation were stable as well (**Fig. 7c-d**). In one exceptional case we found an unexpected decrease in correlation of a specific channel (Day 2 for forward model; Day 16 for inverse model) due to the amplitude change of the measurements. Overall, the predicted neural activities were still highly correlated with the empirical recordings (coefficient = 0.55 and 0.50 for forward and inverse model).

We then tested the stability in phase analysis based on the percentage of predictions located in each zone of scatter plot. Forward-NBGNet-inferred screw ECoG were highly phase-synchronized with the real recordings across different days (**Fig. 7e**). More predictions were always located in the strong synchrony zone than in the poor one (+39%, +21%, +23%, +12%, +23% for Day 1, 2, 4, 12, 16, respectively). Also, reconstructed LFPs were consistently in sync (**Fig. 7f**). Although smaller portion of good synchrony was observed in Day 2 (−4%) and Day 12 (−10%), more synchronous predictions were inferred across sessions (+4%/session).

The NBGNet maintained the capability of reconstructing latent dynamics during the repeated movement generation for the full length of recordings from the monkey (**Fig. 7g,h**). The stability held for a range of manifold dimensionalities from 1 to 3. As we found in Day 1 (**Fig. 3e,i**), descending trend in the correlations of neural modes was observed for multiple days. The average temporal correlations also showed similar results for both forward and inverse model. We then tested whether NBGNet inferences could be used to predict behavior in different sessions. It was noted that the classifiers performed as well as that trained in different sessions (**Fig. 7i**). These results provided evidence that NBGNet-derived signals can be employed to predict behavioral variables with similar accuracy as compared with the ground-truth signals for multiple sessions.

### Comparison of NBGNet and well-known algorithms

We compared the NBGNet with two conventional alternative approaches, specifically a sphere head model^25,47^ and RNN (**Supplementary Fig. 9-12**). Assuming a multi-layered spherical head where each layer represents each brain tissue, the sphere head model provides analytical formulas describing the contribution from current sources to EEG potentials, which are further utilized to approximate the inverse computation. We also applied the machine learning technique, RNN, as a purely date-driven alternative. NBGNet outperformed the purely data-driven RNN and electrophysiology-based sphere head model for multiple days (**Fig. 7j**). As expected, the performance of analytical solution was the worst due to the non-high-density recordings and the unrealistic assumptions (e.g., isotropic conductivity of the medium). While RNN clearly performed better than the sphere head model, it was not competitive enough as compared with the NBGNet. In this demonstration, the NBGNet gave a more accurate inference consistently over multiple days.

## Discussion

The brain consists of a hierarchical system with multiple levels of organization^48^. To gain a better understanding of brain computations, interest in multi-scale interaction among the genetic, cellular, and macroscale levels has been growing recently, inspiring a shift from emphasizing neural communication in individual scales to exploring the potential associations between scales. However, how these scales are interlinked is still an open question. NBGNet addresses an unmet need to capture the implicit relations of multi-scale brain activity. In the present study we have shown that the neural activity at one scale can be inferred from one another and the performance remains stable over multiple days. The underlying dynamics embedded in NBGNet illustrate how the activity at one scale communicate with other scales, which is a key factor in uncovering the mechanistic understanding of brain computations and the mediation of the behaviors.

Assuming the brain computation as a nonlinear operator, we employ the deep learning technique to approximate the nonlinear mapping in the BG-derived dynamical systems which describe the translations between different scales. Although channel selection is based on the spatial relationships, NBGNet’s performance is not dependent upon the depth (**Supplementary Fig. 13**), but on the regions (**Fig. 2**). Forward-NBGNet captures the internal dynamics for performing a center-out task and thus accurately reconstructs the task-related neural activity in premotor, prefrontal cortex, and primary motor cortex. Similarly, inverse-NBGNet-inferred activity matches the ground truth at motor area other than the primary somatosensory cortex. As inverse model is developed by nonlinearizing the inversion of linear forward mapping rather than the direct inversion of nonlinear forward mapping, a slightly poorer performance is observed at more ventral brain region. Since the channel-to-channel communication is modeled individually, scaling up to more channels is expected to mitigate the unexpected mismatch. Evidenced with the failure of capturing the noise from unstable recordings, dynamics embedded in the NBGNet are useful for disambiguating brain computations.

The guiding factor in model evaluation is utilizing comprehensive metrics. This is especially important for neuroscientific research. A perfect performance in one metric may not guarantee the same observation in another. RMSE (**Fig. 1, 6**) is used to indicate the absolute measure of fit. Because correlation and coherence are the measures of similarity analyzed in time- and phase-domain, we also employ cross-correlation (**Fig. 2**) and phase synchrony (**Fig. 3**) for assessment. Capability of reconstructing the low-dimensional latent dynamics is essential as it holds a key to the understanding of neural mechanisms (**Fig. 5**). Additionally, we consider the decoding accuracy as an indicator for the applicability of brain-machine interface (**Fig. 6**). Finally, the predictive power without retraining the model over a long period has recently drawn much attention for the field of neural engineering (**Fig. 7**). These analyses are commonly used in neuroscience research. We validate the NBGNet as a reliable approach with these comprehensive metrics, thus showing a broad applicability in this field.

The bias-variance tradeoff is a critical problem in statistics and machine learning, where the simple models have a lower variance yet a higher bias, and the complexity of the model can reduce the bias but increase the variance. It is thus expected that the NBGNet outperforms the analytical sphere head model and the RNN. With the assumptions of the signal sources and the conductivity of the brain tissues, the sphere head model provides a simple solution, but it leads to a large bias error. The data-driven RNN enables the approximation of nonlinear dynamics; however, a large variance, or the so-called “overfitting,” can be observed. Therefore, to make a fair comparison, we trained the RNN with appropriate regularization. However, the regularized RNN is still a black box without any clue of connections in the model. Combining both electrophysiological modeling and deep learning technique, NBGNet, with the complexity lying between sphere head model and RNN, can capture the patterns in the training data and adapt itself to unseen data, holding a great potential to resolving the bias-variance dilemma.

NBGNet will be helpful in investigating the underlying dynamics in multi-scale brain networks. Modeling the neural activity at disparate scales yields the causal interactions among multiple levels, which is crucial in illuminating the mechanistic understanding of brain computation. Effective connectivity extracted from the NBGNet exhibited both unique and shared patterns of both visual feedback and voluntary movement (**Fig. 4**), suggesting that the NBGNet serves a powerful tool to study the brain computation. Whereas current work focuses on cross-scale interaction, within-scale communication can be incorporated for comprehensive modeling. Additionally, NBGNet can potentially improve the applicability of brain-machine interfaces by inferring the brain activity with increased signal-to-noise ratio and even combining multi-scale activity^49^. Moreover, the inverse computation to reconstruct the activity at the uncovered brain regions makes LFP-derived whole-brain dynamics available. Taken together, our work represents an important step forward towards the mechanistic modeling of multi-scale neural activity, which may facilitate our understanding of neuropathological activity and the development of clinical devices and rehabilitative therapies to treat abnormal neural activity underlying dysfunctional behaviors.

## Data availability

All neural data in this study are available from the corresponding author upon reasonable request.

## Code availability

Python scripts for the model will be made available on GitHub.

## Acknowledgements

We thank José del R. Millán from Clinical Neuroprosthetics and Brain Interaction (CNBI) lab at University of Texas at Austin for extensive discussion and suggestions. This research was partially funded by the Defense Advance Research Project Agency (DARPA) under cooperative agreement W911NF-14-2-0043 issued by the Army Research Office contracting office in support of DARPA’s SUBNETS program (to J.M.C.). H.-C.Y. is supported in part by National Institutes of Health (GM129617) and Robert A. Welch Foundation (F-1833). Y.-I.C. is supported by the University Graduate Continuing Fellowship at UT Austin.

## Contributions

Y.-J.C. and S.R.S. conceived the project, performed data analysis, and wrote the manuscript with input from all of the authors. J.M.C. and S.R.S. designed and performed the experiments. Y.-J.C. and Y.-I.C. developed the algorithmic approach. Y.-I.C., H.-C.Y., J.M.C., and S.R.S. advised data analysis.

## Methods

### The NBGNet model

#### The NeuroBondGraph Network

The NBGNet model is a sparse recurrent neural network (RNN) that captures the system dynamics functionality, especially causality. RNNs make use of sequential information, assuming the output being depended on the previous computations. Accordingly, RNNs have the memory to capture the information about what has been calculated so far. Evidenced with the fact that the recurrent connections exist in visual cortex and hippocampus^50^, RNN is crucial to capture the neural dynamics. Unlike a “black box” model, RNNs in NBGNet are sparsely connected, creating a “gray-box” recurrent neural model using a priori knowledge about the system from the Bond Graphs (BGs)^20^. BGs are regarded as powerful tools to model the physical system by investigating how energy is generated, transferred, transformed, and dissipated among the different system components. BGs allow us to obtain the full representation of the dynamics of the system, and then extract the nonlinear dynamic equation describing the system. Starting with modeling the signal translation between brain activity recorded from different modalities, we can obtain the underlying nonlinear dynamic equations and generate the corresponding NBGNet. In the end, the well-trained network served as the solution to inferring the neural activity from one another.

#### Bond Graphs modeling

Bond graph is a graphical representation of a physical system that allows us to describe the system with a state-space representation. BG consists of the “bonds,” representing the power which is categorized into two parts: flow and effort, and the elements, including single-port, and multi-port components. Literally, single-port elements only have one port for the linkage. Such elements include (1) sources, denoted as *S*_*e*_ or *S*_*f*_, serving as the input effort or flow for the system, (2) sinks, denoted as the same as the sources, serving as the output effort or flow for the system, (3) inertia elements, denoted as *I*, are the ones that store energy, (4) resistance elements, denoted as *R*, are the elements that dissipate energy, and (5) compliance elements, denoted as *C*, are the elements that store potential energy. Furthermore, multi-port elements such as 0-junctions and 1-junctions are employed to split power across their ports. The properties of 0-junctions are that all efforts are equal across the bonds and the sum of flow in equals to the sum of flow out. In contrast, 1-junctions represent that all flows are equal across the bonds and the sum of effort in equals to the sum of effort out. The main concept of BG is to link each component in the graph with the “bonds” which represent the bi-directional exchange of energy. Each bond has two features, half-arrow and causality. Half-arrow is used to provide the sign convention for the work being done. Accordingly, sources will always have the arrow pointing away from the element, while the rest of the single-port elements will have the arrow pointing into the elements. Causality in BG denotes which side of the bond governs the instantaneous effort and flow. The causal stroke is assigned to one end of the bond to represent the side with causality defines the flow while the opposite end defines the effort. In general, since two passive components, *I* and *C*, exhibit time-dependence behavior, they have preferred causal orientation with *C* component defining the effort and *I* component defining the flow. Since BGs are domain neutral, where energy in different domains (e.g., linear mechanical, angular mechanical, electromagnetic, and hydraulic) can be transferred into each other with a constant, this dynamical modeling can incorporate physical systems in multiple domains.

#### Bond Graphs forward modeling

The interactions between recording systems were modeled based on the physiology of brain tissue and its effect on the electrical signal flow. Screw ECoG signals were recorded within the skull while LFP signals were measured within the cortex. Therefore, the brain tissues between the recording location of screw ECoG and LFP are skull, dura mater, and cortex. Assuming that the LFP recordings are the electrical source, the screw ECoG is the measurement of the voltage, and brain tissues represented the effective impedance, we can model it as an electrical circuit. The skull is the bony structure which contains sinus cavities and numerous foramina. From the anatomy, the skull consists of three layers: a spongy bone layer in the middle of two compact bone layers. The cavities in the spongy bone can be modeled as a capacitance that provides a potential inside them. The compact bone and the trabeculae of the spongy bone can be modeled as resistances. Various potential paths for electrical signals to travel are taken into account to construct a simplified electrical model to study the signal propagation. As dura mater is a thick membrane surrounding the brain, dura model is built with the effective resistance and capacitance in parallel. Although cortex is composed of folded grey matter, it is modeled as an effective resistance to simplify the complexity. Combing together, LFP-screw ECoG transmission electrical circuit can be established (**Supplementary Fig. 1a**). With the effective electrical circuit for electrical signal pathways, Bond Graph is then generated (**Supplementary Fig. 1b**), where the charges, *q*, are represented the states of the physical systems. Given the causality assignment, the system is 3^rd^ order. The ordinary differential equations for the system are derived as follows:

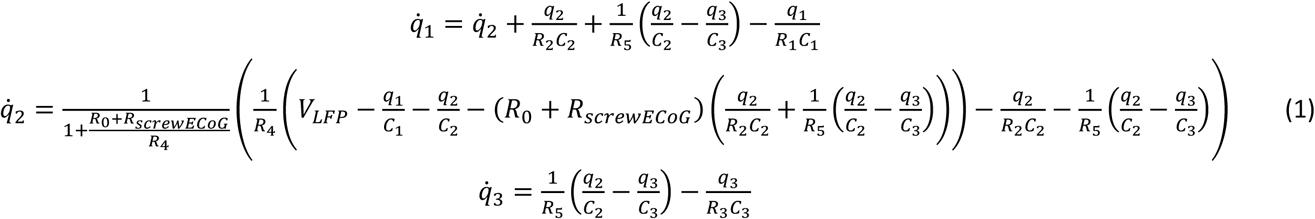

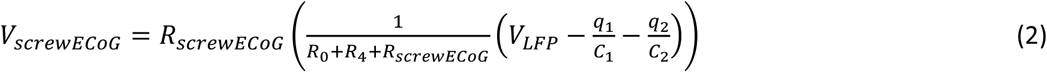

While the equations above represent the ideal condition where the resistance *R* and the capacitance *C* are linear. Considering the uncertainty and continuous changing of human’s brain tissue, nonlinearity is introduced in the equation:

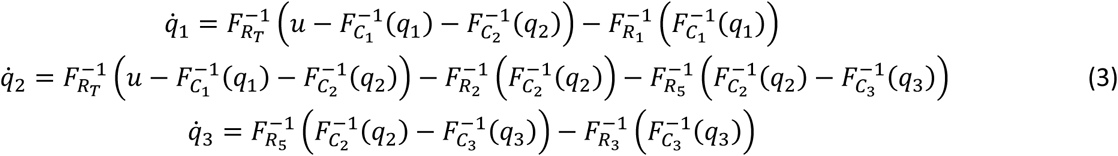

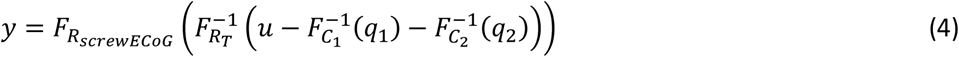

where, *R*_*T*_ represents *R*_0_ + *R*_4_ + *R*_*screwECoG*_, *u* represents *V*_*LEP*_, *y* represents *V*_*screwECoG*_, and *F*(*·*) is a nonlinear function to be determined.

#### Bond Graphs inverse modeling

The multi-variable time varying Bond Graph forward model, equation (1-2), can be expressed as the state-space representation,

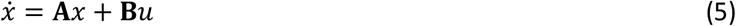

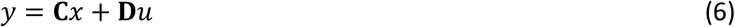

where, *x* = [*q*_1_, *q*_2_, *q*_3_]^*T*^, *u* = *V*_*LEP*_, *y* = *V*_*screwECoG*_,

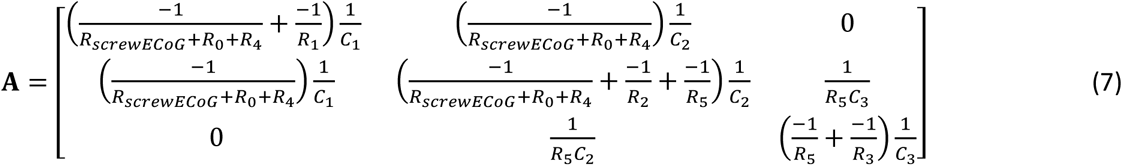

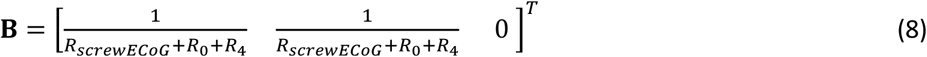

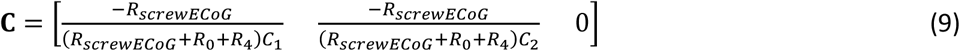

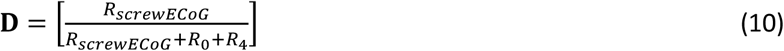

The inversion algorithm for multi-variable system were obtained by Sain and Massey^51^,

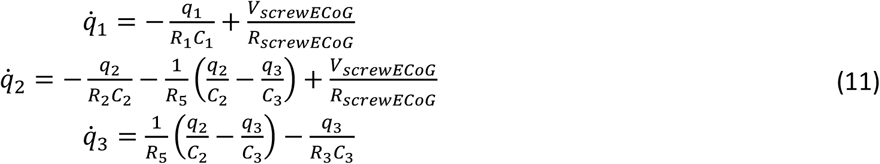

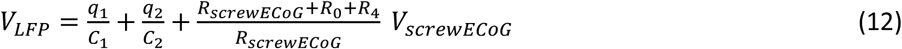

As forward model, nonlinearity is introduced in the equation as well,

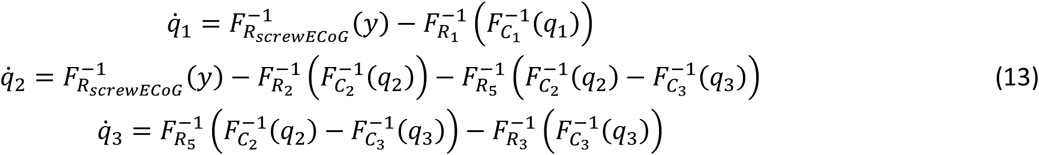

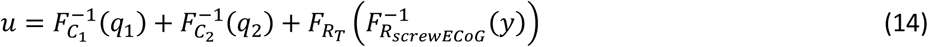

where, *R*_*T*_ represents *R*_0_ + *R*_4_ + *R*_*screwECoG*_, *u* represents *V*_*LEP*_, *y* represents *V*_*screwECoG*_, and *F*(*·*) is a nonlinear function to be determined.

#### NeuroBondGraphs Network modeling

To approximate the unknown nonlinear relationship, artificial neural network technique is introduced to incorporate the knowledge obtained from the pattern recognition capabilities of neural network and the physical information about the system via BG. Each nonlinear function is approximated through a multi-layer perceptron (MLP) unit. The architecture of NBGNet is a sparse recurrent MLP network which can identify the nonlinear dynamics recursively.

#### The full NBGNet inference model

The full NBGNet, both forward and inverse models, are run in the following way. First, a data trial is chosen. Then, for each time step from 1 to *T*, the network is updated and eventually predicts the output according to

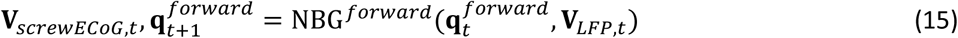

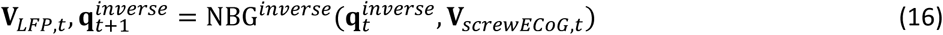

After training, the full model can be run, starting with any single trial or a set of trials corresponding to the particular experimental conditions, to obtain the associated dynamic states, and inferred target signals for that trial or condition.

#### The loss function

To optimize our model, we would like to minimize the mean-squared-error (MSE) of the data, which measures the average of the squares of the errors. Given the ground-truth measurement ***Y***_*gt*_ and the model prediction ***Y***_*pre*_, the MSE is calculated as follows,

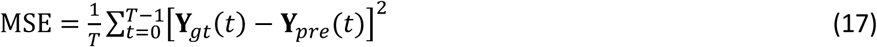

where, *T* is the number of time points in the given trial. The loss function will be minimized by backpropagating the error and update the parameters of the model iteratively.

#### Hyperparameters and further details of NBGNet implementation

The major hyperparameters for forward and inverse model is the number of hidden nodes in the MLP unit for nonlinear mapping estimation and the time step. For both forward and inverse model, 7 nodes were utilized in MLP units, and the time step of NBGNet was equal to the data bin size. To help avoid over-fitting, we selected different trials of data for model training when NBGNet has been updated for twenty times. Furthermore, NBGNet is sparsely connected rather than fully connected, which reduces the complexity of the model.

NBGNets were randomly initialized by Glorot uniform initializer and optimized using adaptive moment estimation (Adam) optimizer with a starting learning rate of 1 × 10^−3^. The framework was implemented with TensorFlow version 1.15.0 and Python version 3.7.4 in the Microsoft Windows 10 operating system. The training was performed on a consumer-built desktop equipped with 4 GPUs (Gigabyte RTX 2080 Ti graphics cards, NVIDIA) and a Core i9-9820X CPU @ 3.3 GHz (Intel). To monitor overfitting, a portion of the data is set aside as the validation set, and these data are not used to update the weights of NBGNet. Instead, they are used to evaluate the predictive performance on held-out data. Here we used a ratio of 9:1 between training and validation data.

### Experimental model and subject details

A male rhesus macaque was used in these experiments. All experiments were approved by the Animal Care and Use Committee at the University of California, Berkeley. The subject was approximately 6 years of age at the time of data collection.

### Behavioral tasks

The macaque monkey was trained in a center-out task. Briefly, the subject was trained to use a joystick to move a cursor on a computer screen from a center target to a peripheral target. The joystick was located in front of the primate chair and the subject was free to use either hand to control the joystick during the experiment. In the task, the subject was trained to fold the cursor at the center target for 320 ms. Then the subject was presented with one of the eight outer targets, equally spaced in a circle and selected randomly with uniform probability. The subject moved the cursor to the peripheral target and held the cursor inside the target for 320 ms. A trial was considered to be successful if the subject completed the 320 ms hold-center followed by holding at the peripheral target for 320 ms. The reward was scheduled after a successful trial, where a custom-programmed Arduino triggered the reward system to deliver a small amount of juice to the subject.

### Scale-dependent analysis

To evaluate how close the model predictions are to the ground-truth signals, root mean square error (RMSE) is commonly used to indicate the absolute fit of the model. RMSE is defined as the square root of the mean of the square of the error,

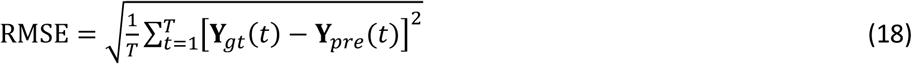

where the ***Y***_*gt*_ represents ground-truth measurement, ***Y***_*pre*_ represents the model prediction, and *T* is the number of time points in the given trial.

### Similarity analysis

Similarity of two time series signals also conveys an important message whether two time series signals exhibit similar shape of oscillation. Here we use Pearson correlation coefficient to measure how highly correlated two time series signals are.

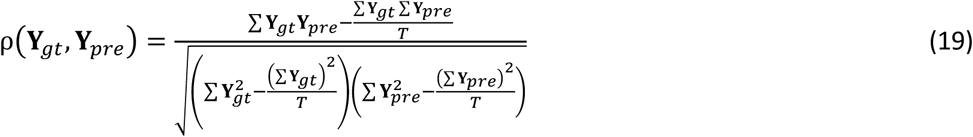

where the ***Y***_*gt*_ represents ground-truth measurement, ***Y***_*pre*_ represents the model prediction, and *T* is the number of time points in the given trial.

### Phase analysis

On top of the time-domain, phase-domain reveals other characteristics that are not visible in time-domain. Especially in neural engineering field, the synchronization of the neural activity is one of the properties that can only be quantified in phase domain. Given a pair of signals, ***s***_1_(*t*) and ***s***_2_(*t*), which have been band-pass filtered to a frequency range of interest, Hilbert transform, ***HT***[·], will be applied to obtain the corresponding analytical signals, ***z***_1_(*t*) and ***z***_2_(*t*),

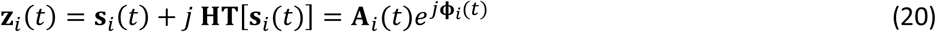

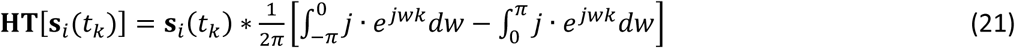

where *k* = 1 to *T*, ***A***_*i*_(*t*) represents the instantaneous amplitude, and ***Φ***_*i*_(*t*) represents the instantaneous phase. In order to obtain a comprehensive view, we utilized two metrics: phase-locking value and phase synchrony index. Phase locking value^30^, *PLV*, or so-called mean phase coherence^31^, is defined as,

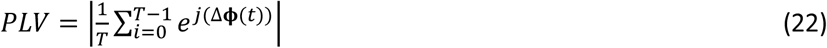

It is utilized to measure the intra-trial variability of the phase difference between two signals. In addition, the phase of phase-locking can be extracted to evaluate the mean phase different across time.

On top of *PLV*, we are also interested in the instantaneous performance, and thus we introduce phase synchrony index. First, provided with the instantaneous phase of two time series signals, ***Φ***_1_(*t*) and ***Φ***_2_(*t*), the instantaneous phase synchrony, *IPS*, which measures the phase similarity at each timepoint, is calculated by

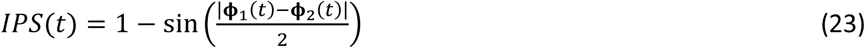

where the phase is in the unit of degree. We define 45° as the threshold of the phase difference. When the phase difference is less than 45°, *IPS* is greater than 0.62. We then calculate the ratio of the time with the *IPS* greater than 0.62, termed phase synchrony index, *PSI* (**Supplementary Fig. 4a**),

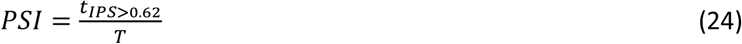

To determine the level of the phase synchrony, we categorize the two-dimensional scatter plot of *PSI* and *PLV* into four sections with both thresholds as 0.5: Zone 1 (low *PSI* and low *PLV*) indicates poor synchronization, Zone 2 (low *PSI* and high *PLV*) indicates medium synchronization, Zone 3 (high *PSI* and low *PLV*) indicates medium synchronization, and Zone 4 (high *PSI* and high *PLV*) indicates perfect synchronization (**Supplementary Fig. 4b**).

### Neural latent dynamic analysis

To characterize the latent dynamics associated with the recorded or reconstructed neural activity in each trial, we analyzed the filtered signals, which were obtained by applying a bandpass filter at 12.5 Hz and 30 Hz, in the window starting at movement onset and ending 600 ms after movement onset. Such a window was selected was due to the interest in movement execution during the trial. For each trial, we obtained the data matrix *D* of dimension *n* by *T*, where *n* is the number of recorded channels, *T* is the number of time points in the given trial. Then we computed the low-dimensional manifold by applying principal component analysis (PCA) to *D*. The resulting PCs are the linear combination of measurements of all the channels. We would then rank these PCs based on the amount of neural variance explained by each PC explains. We keep only three leading PCs to represent the low-dimensional manifold, where these three leading PCs, or called neural modes, explain most of the variance in the data matrix (**Supplementary Fig. 5**).

Assuming the latent dynamics captured by the neural recordings and the reconstructed latent dynamics inferred from NBGNet’s predictions are the projections on the different manifolds from the true latent dynamics^52^, we expect to identify the embedding space that true latent dynamics is located by using canonical correlation analysis (CCA). In CCA, given a pair of two latent trajectories, ***P***_*A*_ and ***P***_*B*_, linear transformations for each trajectory are identified to make the linearly transformed latent trajectories, 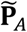 and 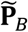, maximally correlated. First, QR decomposition is applied to both latent trajectories,

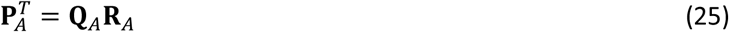

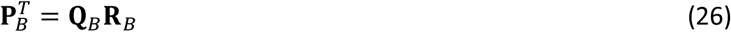

Then the singular value decomposition is performed on the inner product of ***Q***_*A*_ and ***Q***_*B*_.

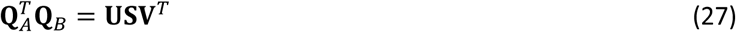

The transformation matrix, **T**_*A*_ and **T**_*B*_, is obtained by,

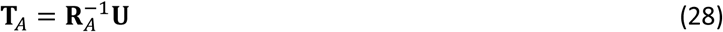

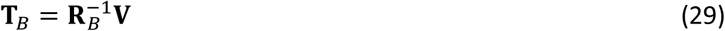

Accordingly, the transformed latent trajectories are given by

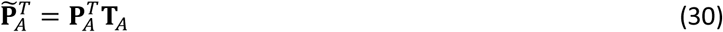

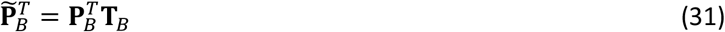

The correlation between the transformed latent trajectories, termed canonical correlation (CC), is obtained by the Pearson correlation coefficient. As CC is sorted from the largest to the smallest in CCA, we expect to observe a descending order from neural mode 1 to mode 3.

### Features selection for decoding the direction of the movement

We introduced several features per channel as candidates for the decoder and selected the leading number of features for further analysis. For each channel, we obtained total 34 features, including root mean square (RMS), mean frequency (MF), waveform length (WL), and the power at certain frequency ranged from 10 – 40 Hz (step size as 1 Hz).

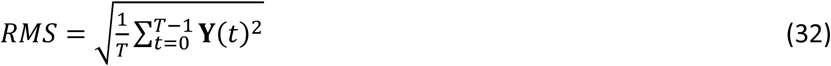

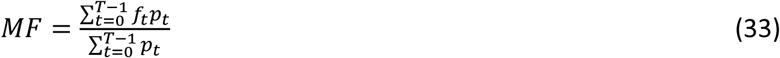

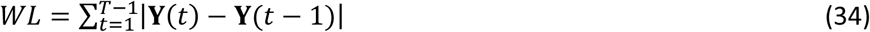

where ***Y***(*t*) represents the neural signals, *T* is the number of time points in the given trial, *f*_*t*_ and *p*_*t*_ are the frequencies of the power spectrum and the corresponding amplitude.

To determine the features to be selected, we calculate the Fisher score for each candidate and select to the leading features. The Fisher score, *F*(*x*^*i*^), for the *i*-th feature, *x*^*i*^, is computed by

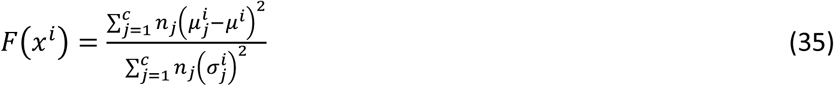

where 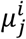 and 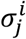 are the mean and standard deviation of the *j*-th class corresponding to the *i*-th feature, *μ*^*i*^ denotes the mean of the whole data set corresponding to the *i*-th feature, *n*_*j*_ represents the size of the *j*-th class, and *c* is the total number of classes. After computing the Fisher score for each feature, we selected the top-six ranked features to predict the subject’s behavior. The features selected for analysis are summarized in **Supplementary Table 3**.

### Predicting target direction from LFP and screw EcoG

To test whether the reconstructed activity from NBGNet maintain movement-related information, we built linear decoders to predict the direction of the movement based on the neural activity. Our hypothesis was that our NBGNet inference should provide as accurate predictions as the neural recordings. To test this hypothesis, we compared the predictive accuracy of seven types of decoders: (1) a decoder trained and tested based on screw ECoG; (2) a decoder trained and tested based on reconstructed screw ECoG inferred by forward-NBGNet; (3) a decoder trained and tested based on LFP; (4) a decoder trained and tested based on reconstructed LFP inferred by inverse-NBGNet; (5) a decoder trained and tested based on screw ECoG and LFP; (6) a decoder trained and tested based on reconstructed screw ECoG and LFP; and (7) a decoder trained and tested based on reconstructed LFP and screw ECoG. All decoders were linear discriminant analysis that used the selected features (**Supplementary Table 3**) as inputs to predict the direction of cursor’s movement. They were trained and tested on the same day, using nine-fold cross-validation procedure to protect against overfitting. Chance-level performance was obtained by shuffling the dataset. As expected, all the predictive accuracy was higher than the chance-level, which was around 12.5 %.

### Comparison methods

To evaluate performance of NBGNet as compared to other existing algorithms, we implemented two approaches from different perspectives: electrophysiology-based sphere head model and data-driven recurrent neural network.

#### Sphere head model

Sphere head model is widely used to either compute the contribution from the current dipoles to the electrical potentials recorded at scalp electroencephalography (EEG) or estimate the current dipole sources based on the scalp potentials. Assume the head can be modeled as a four-layered sphere where each layer represents each tissue: brain, cerebrospinal fluid, skull, and scalp, respectively. Using the quasi-static approximation of Maxwell’s equations and the volume-conductor theory, the electrical potential, **Φ**(**r**, *t*), can be calculated by the following Poisson equation^53^:

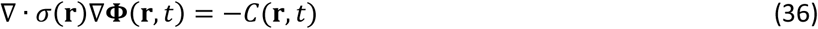

where *σ*(**r**) represents the position-dependent conductivity of the medium, and *C*(**r**, *t*) is the density of the current sources. Assuming the conductivity to be isotropic, the solution provided by sphere head model to equation (36) is

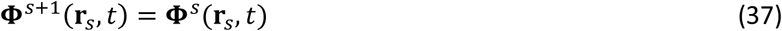

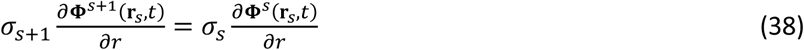

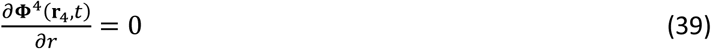

where each layer is labeled by *s* = 1 to 4. Here we assume the dipole is radial with the magnitude *p*(*t*) at location *r*_*z*_, the analytical solution to equation (37 – 39) is given by,

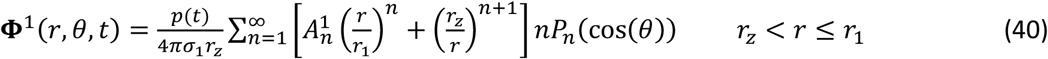

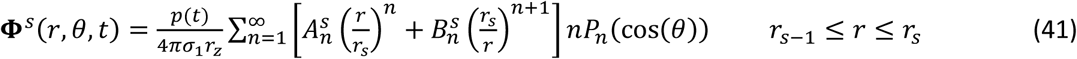

where **Φ**^*s*^(*r, θ, t*) is the extracellular potential measured at radius *r* and the angle *θ* between measurement and dipole location vectors in the shell *s, r*_*s*_ represents the radius of sphere *s*, 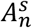 and 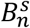 are the coefficients depending on the radius and conductivities of each medium, where are defined in^16^, and *P*_*n*_(cos(*θ*)) represents the *n*-th Legendre Polynomial. As the solution is implemented into the case where we have *n*_*d*_ current dipoles and *n*_*r*_ recording units, a linear transformation matrix *F* of dimension *n*_*r*_ by *n*_*d*_ can be obtained and utilized to convert the dipole moment vectors **X** into the electrical potential ***Y***, given by ***Y*** = *F***X**. This is so-called forward mapping. When we performed inverse mapping to estimate **X** from ***Y***, we need to solve such an underdetermined system with pseudo-inverse by minimizing the following equation,

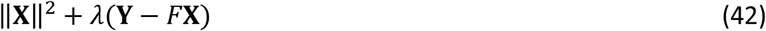

The solution to minimizing the equation (42) is given by,

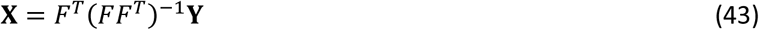

Here we segment the brain into 3600 segments (*n*_*d*_ = 3600), where each segment includes a potential current dipole source. Since our data for comparison does not include dipole sources, we adapt the algorithms into two step computation for both forward and inverse model. In forward model, we performed inverse mapping from LFP toward estimated dipole sources followed by a forward mapping from the estimated dipole sources toward screw ECoG recordings. Similarly, in inverse model, we performed inverse mapping from screw ECoG toward estimated dipole sources followed by a forward mapping from the estimated dipole sources toward LFP recordings. The parameters utilized are summarized in **Supplementary Table 4**.

#### Recurrent neural network (RNN)

RNN is a commonly-used deep learning method to model a nonlinear dynamical system which includes three characteristics: nonlinearity, recurrent connection, and hidden dynamic states. In order to handle the long-term dependency embedded in the neural activity, Long Short-Term Memory (LSTM)^54^ is often implemented, where in each time point, model can determine the information to be forgotten, to be stored, or new information to be added. LSTM-based RNN utilized in this work for comparison consists of two LSTM layers with 64 and 32 units, two hidden layers with 64 and 16 nodes, and the output layer. To avoid overfitting, we trained the RNN with L2 regularization and dropout. The relevant hyperparameters were optimized via Bayesian optimization. The training details, including training iteration, the split ratio of training and validation data, and the choice of optimizer, were set to be the same as NBGNet to ensure a fair comparison.

## Data availability

The datasets analyzed for this manuscript will be shared upon reasonable request.

## Code availability

All analyses were implemented using custom Python code. Code to replicate the main results will be shared upon reasonable request.

## Reference

1. Churchland, M. M., Santhanam, G. & Shenoy, K. V. Preparatory Activity in Premotor and Motor Cortex Reflects the Speed of the Upcoming Reach. Journal of Neurophysiology 96, 3130–3146 (2006).

2. Churchland, M. M., Cunningham, J. P., Kaufman, M. T., Ryu, S. I. & Shenoy, K. V. Cortical Preparatory Activity: Representation of Movement or First Cog in a Dynamical Machine? Neuron 68, 387–400 (2010).

3. Vyas, S., O’Shea, D. J., Ryu, S. I. & Shenoy, K. V. Causal Role of Motor Preparation during Error-Driven Learning. Neuron 106, 329-339.e4 (2020).

4. Mauk, M. D. & Buonomano, D. V. The Neural Basis of Temporal Processing. Annual Review of Neuroscience 27, 307–340 (2004).

5. Remington, E. D., Egger, S. W., Narain, D., Wang, J. & Jazayeri, M. A Dynamical Systems Perspective on Flexible Motor Timing. Trends in Cognitive Sciences 22, 938–952 (2018).

6. Chaisangmongkon, W., Swaminathan, S. K., Freedman, D. J. & Wang, X.-J. Computing by Robust Transience: How the Fronto-Parietal Network Performs Sequential, Category-Based Decisions. Neuron 93, 1504-1517.e4 (2017).

7. Chaudhuri, R. & Fiete, I. Computational principles of memory. Nature Neuroscience 19, 394–403 (2016).

8. Miller, E. K., Lundqvist, M. & Bastos, A. M. Working Memory 2.0. Neuron 100, 463–475 (2018).

9. Macke, J. H., Buesing, L. & Sahani, M. Estimating state and parameters in state space models of spike trains. in Advanced State Space Methods for Neural and Clinical Data (ed. Chen, Z.) 137–159 (Cambridge University Press, 2015). doi:10.1017/CBO9781139941433.007.

10. Byron, M. Y. et al. Gaussian-Process Factor Analysis for Low-Dimensional Single-Trial Analysis of Neural Population Activity. Journal of Neurophysiology 102, 614–635 (2009).

11. Pandarinath, C. et al. Inferring single-trial neural population dynamics using sequential auto-encoders. Nature Methods 15, 805–815 (2018).

12. Buschman, T. J. & Kastner, S. From behavior to neural dynamics: An integrated theory of attention. Neuron 88, 127–144 (2015).

13. Harbecke, J. The methodological role of mechanistic-computational models in cognitive science. Synthese 1–23 (2020) doi:10.1007/s11229-020-02568-5.

14. Lu, H.-Y. et al. Multi-scale neural decoding and analysis. J. Neural Eng. 18, 045013 (2021).

15. Canolty, R. T., Ganguly, K. & Carmena, J. M. Task-Dependent Changes in Cross-Level Coupling between Single Neurons and Oscillatory Activity in Multiscale Networks. PLOS Computational Biology 8, e1002809 (2012).

16. Næss, S. et al. Corrected Four-Sphere Head Model for EEG Signals. Front. Hum. Neurosci. 11, 490 (2017).

17. Michel, C. M. & Brunet, D. EEG Source Imaging: A Practical Review of the Analysis Steps. Front. Neurol. 10, (2019).

18. Vorwerk, J., Aydin, Ü., Wolters, C. H. & Butson, C. R. Influence of Head Tissue Conductivity Uncertainties on EEG Dipole Reconstruction. Front. Neurosci. 13, 531 (2019).

19. Fernández, B., Prabhudesai, A. V., Murty, V. V., Gupta, R. & Chang, W. R. Neurobondgraphs: modeling environment of nonlinear dynamic systems using neural networks and bond graphs. in 75–90 (ASME, 1992).

20. Paynter, H. M. Analysis and design of engineering systems. (MIT Press, 1961).

21. Yeager, J. D., Phillips, D. J., Rector, D. M. & Bahr, D. F. Characterization of flexible ECoG electrode arrays for chronic recording in awake rats. Journal of Neuroscience Methods 173, 279–285 (2008).

22. Choi, H. et al. Long-term evaluation and feasibility study of the insulated screw electrode for ECoG recording. Journal of Neuroscience Methods 308, 261–268 (2018).

23. Merel, J., Botvinick, M. & Wayne, G. Hierarchical motor control in mammals and machines. Nat Commun 10, 5489 (2019).

24. Bond Graphs for Modelling, Control and Fault Diagnosis of Engineering Systems. (Berlin: Springer, 2017). doi:10.1007/978-3-319-47434-2.

25. Grech, R. et al. Review on solving the inverse problem in EEG source analysis. Journal of NeuroEngineering and Rehabilitation 5, 25 (2008).

26. Sanes, J. N. & Donoghue, J. P. Oscillations in local field potentials of the primate motor cortex during voluntary movement. PNAS 90, 4470–4474 (1993).

27. Khanna, P. & Carmena, J. M. Beta band oscillations in motor cortex reflect neural population signals that delay movement onset. ELife 6, e24573 (2017).

28. Dancey, C. P. & Reidy, J. Statistics without maths for psychology. (Pearson education, 2007).

29. Akoglu, H. User’s guide to correlation coefficients. Turkish Journal of Emergency Medicine 18, 91–93 (2018).

30. Lachaux, J.-P., Rodriguez, E., Martinerie, J. & Varela, F. J. Measuring phase synchrony in brain signals. Human Brain Mapping 8, 194–208 (1999).

31. Mormann, F., Lehnertz, K., David, P. & Elger, C. E. Mean phase coherence as a measure for phase synchronization and its application to the EEG of epilepsy patients. Physica D: Nonlinear Phenomena 144, 358–369 (2000).

32. Battaglia, D., Witt, A., Wolf, F. & Geisel, T. Dynamic Effective Connectivity of Inter-Areal Brain Circuits. PLOS Computational Biology 8, e1002438 (2012).

33. Friston, K. J. Functional and Effective Connectivity: A Review. Brain Connectivity 1, 13–36 (2011).

34. Funahashi, S., Bruce, C. J. & Goldman-Rakic, P. S. Mnemonic coding of visual space in the monkey’s dorsolateral prefrontal cortex. Journal of Neurophysiology 61, 331–349 (1989).

35. Arce-McShane, F. I., Ross, C. F., Takahashi, K., Sessle, B. J. & Hatsopoulos, N. G. Primary motor and sensory cortical areas communicate via spatiotemporally coordinated networks at multiple frequencies. PNAS 113, 5083–5088 (2016).

36. Afshar, A. et al. Single-Trial Neural Correlates of Arm Movement Preparation. Neuron 71, 555–564 (2011).

37. Churchland, M. M. et al. Neural population dynamics during reaching. Nature 487, 51–56 (2012).

38. Kaufman, M. T., Churchland, M. M., Ryu, S. I. & Shenoy, K. V. Cortical activity in the null space: permitting preparation without movement. Nature Neuroscience 17, 440–448 (2014).

39. Mante, V., Sussillo, D., Shenoy, K. V. & Newsome, W. T. Context-dependent computation by recurrent dynamics in prefrontal cortex. Nature 503, 78–84 (2013).

40. Sadtler, P. T. et al. Neural constraints on learning. Nature 512, 423–426 (2014).

41. Wold, S., Esbensen, K. & Geladi, P. Principal component analysis. Chemometrics and Intelligent Laboratory Systems 2, 37–52 (1987).

42. Gallego, J. A., Perich, M. G., Miller, L. E. & Solla, S. A. Neural Manifolds for the Control of Movement. Neuron 94, 978–984 (2017).

43. Thompson, B. Canonical Correlation Analysis. in Encyclopedia of Statistics in Behavioral Science (eds. Everitt, B. S. & Howell, D.) (Wiley, 2005). doi:10.1002/0470013192.bsa068.

44. Gallego, J. A. et al. Cortical population activity within a preserved neural manifold underlies multiple motor behaviors. Nature Communications 9, 1–13 (2018).

45. Winkler, A. M., Renaud, O., Smith, S. M. & Nichols, T. E. Permutation Inference for Canonical Correlation Analysis. arXiv reprint 2002.10046 (2020).

46. Duda, R. O., Hart, P. E. & Stork, D. G. Pattern Classification. 2nd Edition. (Hoboken: Wiley, 2000).

47. Hallez, H. et al. Review on solving the forward problem in EEG source analysis. Journal of NeuroEngineering and Rehabilitation 4, 46 (2007).

48. Heuvel, M. P. van den, Scholtens, L. H. & Kahn, R. S. Multiscale Neuroscience of Psychiatric Disorders. Biological Psychiatry 86, 512–522 (2019).

49. Abbaspourazad, H., Choudhury, M., Wong, Y. T., Pesaran, B. & Shanechi, M. M. Multiscale low-dimensional motor cortical state dynamics predict naturalistic reach-and-grasp behavior. Nat Commun 12, 607 (2021).

## Reference

50. Káli, S. & Dayan, P. The Involvement of Recurrent Connections in Area CA3 in Establishing the Properties of Place Fields: a Model. J Neurosci 20, 7463–7477 (2000).

51. Sain, M. & Massey, J. Invertibility of linear time-invariant dynamical systems. IEEE Transactions on Automatic Control 14, 141–149 (1969).

52. Gallego, J. A., Perich, M. G., Chowdhury, R. H., Solla, S. A. & Miller, L. E. Long-term stability of cortical population dynamics underlying consistent behavior. Nature Neuroscience 23, 260–270 (2020).

53. Nunez, P. L. & Srinivasan, R. Electric Fields of the Brain: The neurophysics of EEG. Electric Fields of the Brain (Oxford University Press, USA, 2006).

54. Hochreiter, S. & Schmidhuber, J. Long Short-Term Memory. Neural Computation 9, 1735–1780 (1997).

