## Supplementary Information for "Inferring system-level brain communication through multi-scale neural activity"

**Supplementary Figures:**
Supplementary Fig. 1: Development of the NBGNet
Supplementary Fig. 2: Power spectral density and time-frequency spectrogram for screw ECoG and LFP are demonstrated informative in beta band
Supplementary Fig. 3: Stability of NBGNet's performance over time is evaluated with RMSE
Supplementary Fig. 4: Phase analysis
Supplementary Fig. 5: Identification of neural modes using principal component analysis
Supplementary Fig. 6: Stability of forward-NBGNet's performance over time is evaluated with cross-correlation
Supplementary Fig. 7: Stability of inverse-NBGNet's performance over time is evaluated with cross-correlation
Supplementary Fig. 8: Stability of NBGNet's performance over time is evaluated with phase synchrony
Supplementary Fig. 9: The performance of NBGNet, sphere head model and RNN was evaluated by RMSE
Supplementary Fig. 10: The performance of NBGNet, sphere head model and RNN was evaluated by cross-correlation
Supplementary Fig. 11: The performance of NBGNet, sphere head model and RNN was evaluated by phase locking value
Supplementary Fig. 12: The performance of NBGNet, sphere head model and RNN was evaluated by phase synchrony index
Supplementary Fig. 13: NBGNet's performance is depth-independent

**Supplementary Tables:**
Supplementary Table 1: Screw ECoG electrode layout
Supplementary Table 2: LFP electrode layout
Supplementary Table 3: Selected features for movement behavior decoding
Supplementary Table 4: Parameters for sphere head model

**Supplementary Discussion**

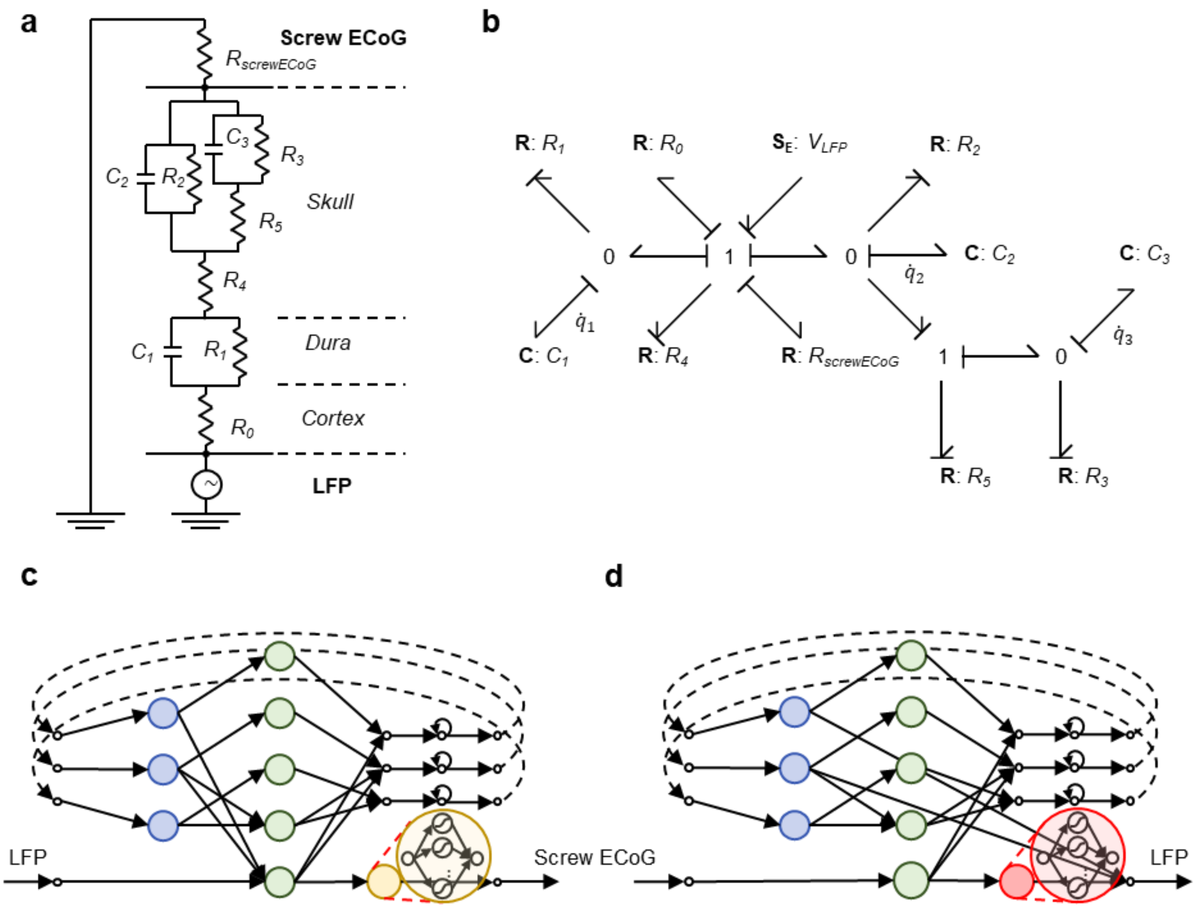


**Supplementary Fig. 1: Development of the NBGNet.** (**a**) LFP-screw ECoG transmission electrical circuit was established based on the effective electrical signal pathway. (**b**)Bond Graph of the physical system illustrated in **a**. (**c-d**) Both forward- and inverse-NBGNet are derived from the system dynamics equations for the LFP-screw ECoG transmission model. (**c**) Schematic of forward-NBGNet architecture, where the colored circles represent a multi-layer perceptron unit. (**d**) Same as **c** for inverse-NBGNet architecture

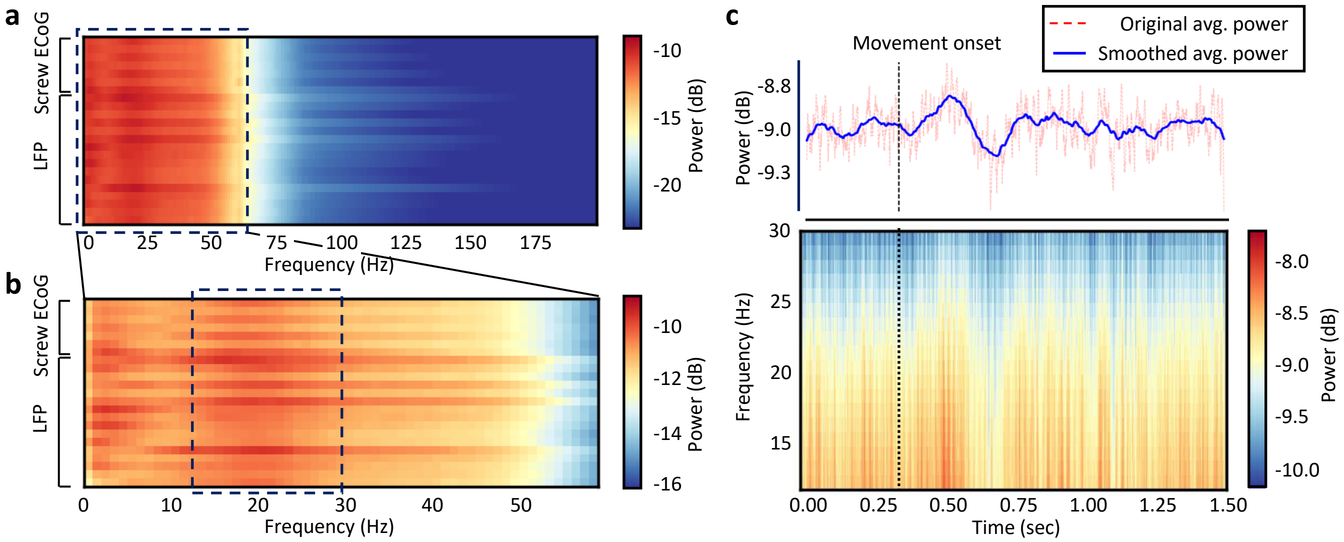


**Supplementary Fig. 2: Power spectral density and time-frequency spectrogram for screw ECoG and LFP are demonstrated informative in beta band.** (**a**) Heatmap of power spectral density (0 – 200 Hz) for screw ECoG and LFP channels. (**b**) Heatmap of power spectral density (0 – 60 Hz, ROI in **a**) for screw ECoG and LFP channels. (**c**) Two-dimensional plot of the spectrogram for screw ECoG channel 4 (lPFC; bottom). Temporal power trajectory is obtained by averaging the power across the frequency, followed by smoothing with 200 time points (top). Dashed line, movement onset.

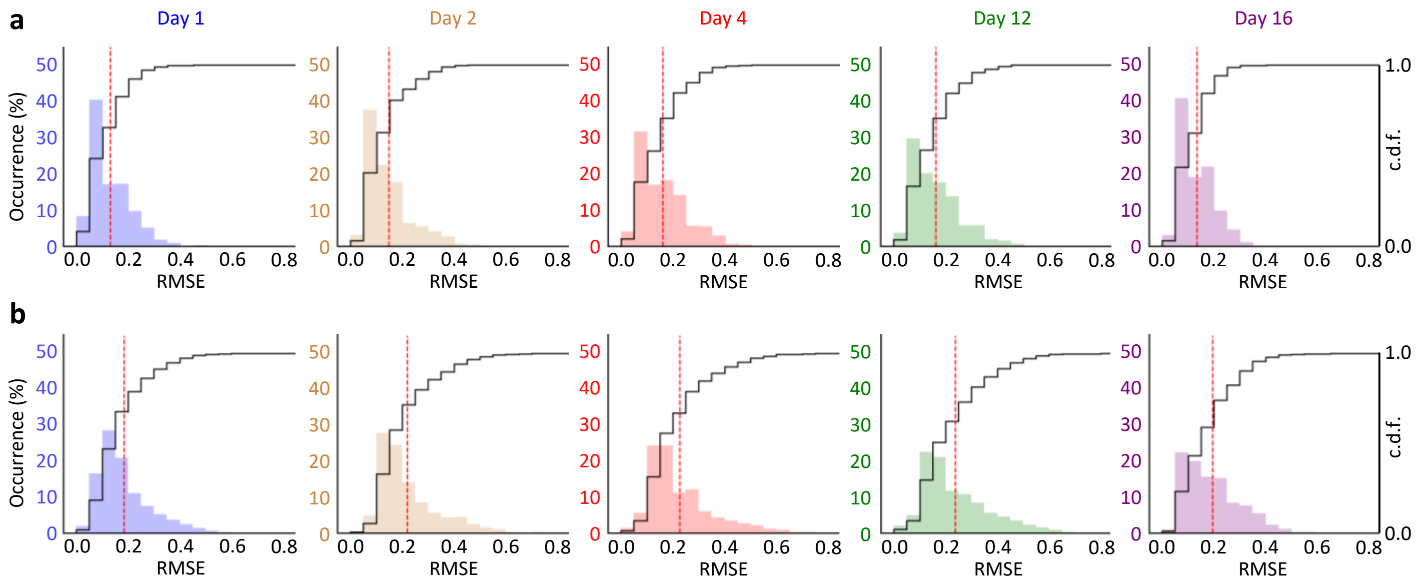


**Supplementary Fig. 3: Stability of NBGNet's performance over time is evaluated with RMSE.** (**a**) Root mean squared error (RMSE) between forward-NBGNet's inference and the ground truth screw ECoG across days. Histogram and cumulative distribution function of RMSE in beta band is close to 0.1 ($10^{-4}$V), indicating that model predictions are close to the ground truths. (**b**) RMSE between inverse-NBGNet's inference and the ground truth LFP across days. Histogram and cumulative distribution function of RMSE in beta band is close to 0.2 ($10^{-4}$V), indicating that model predictions are close to the ground truths as well. (Red dashed line: mean).

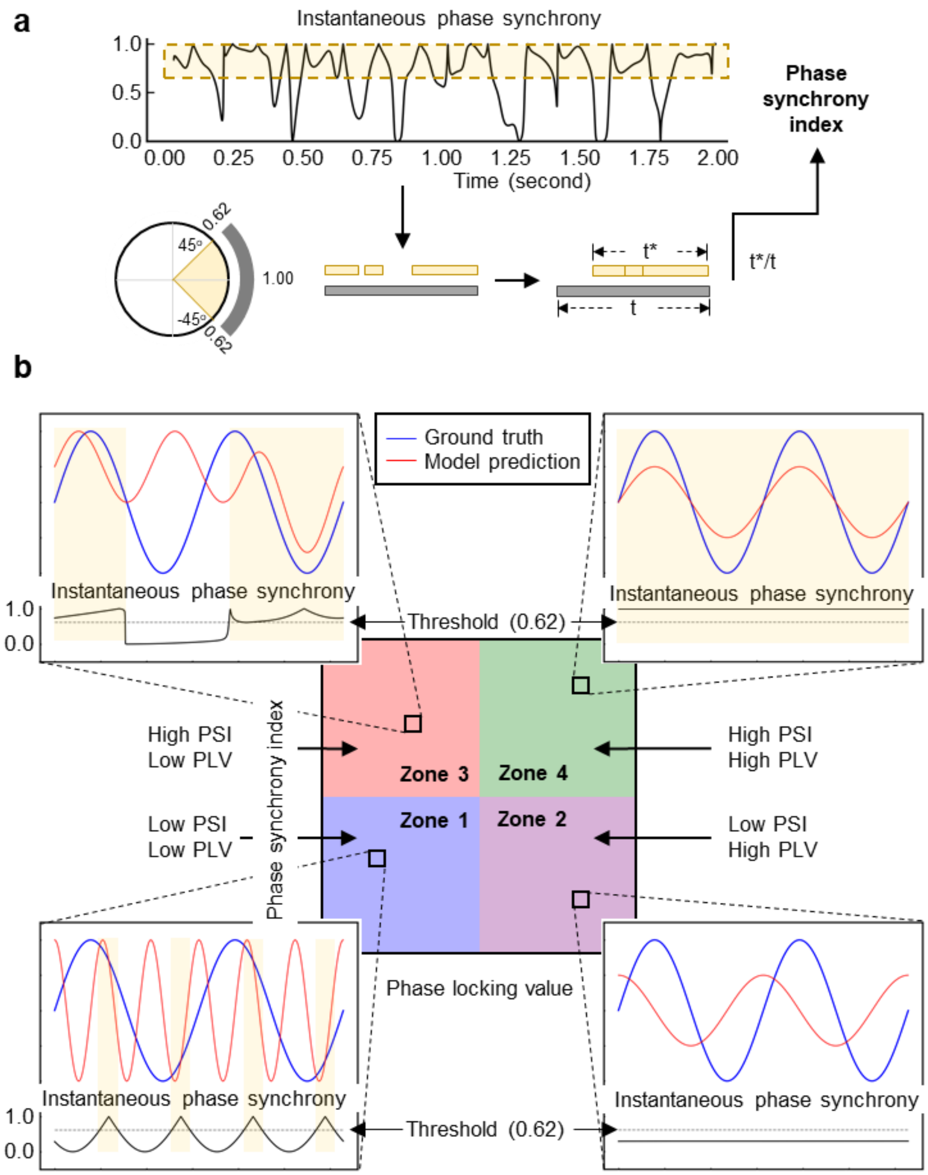


**Supplementary Fig. 4: Phase analysis.** (**a**) Phase synchrony index quantifies the percentage of the time for two time series signals exhibiting high degree of synchrony. Instantaneous phase synchrony is first calculated from the phase of two time series signals. Strong phase similarity is indicated as the instantaneous phase synchrony is greater than 0.62; in other words, the instantaneous phase difference between two signals is less than 45°. The phase synchrony index is then obtained by dividing the time segments meeting the criteria (yellow) by the total time. (**b**) Phase synchrony plot characterizes the details of phase similarities. Two-dimensional scatter plot of phase synchrony index and phase locking value can be categorized into 4 sections with both thresholds as 0.5: Zone 1, poor synchronization; Zone 2 and 3, moderate synchronization; Zone 4, perfect synchronization. Simulated data (top) and the corresponding instantaneous phase synchrony (bottom) indicate the representative cases in each zone.

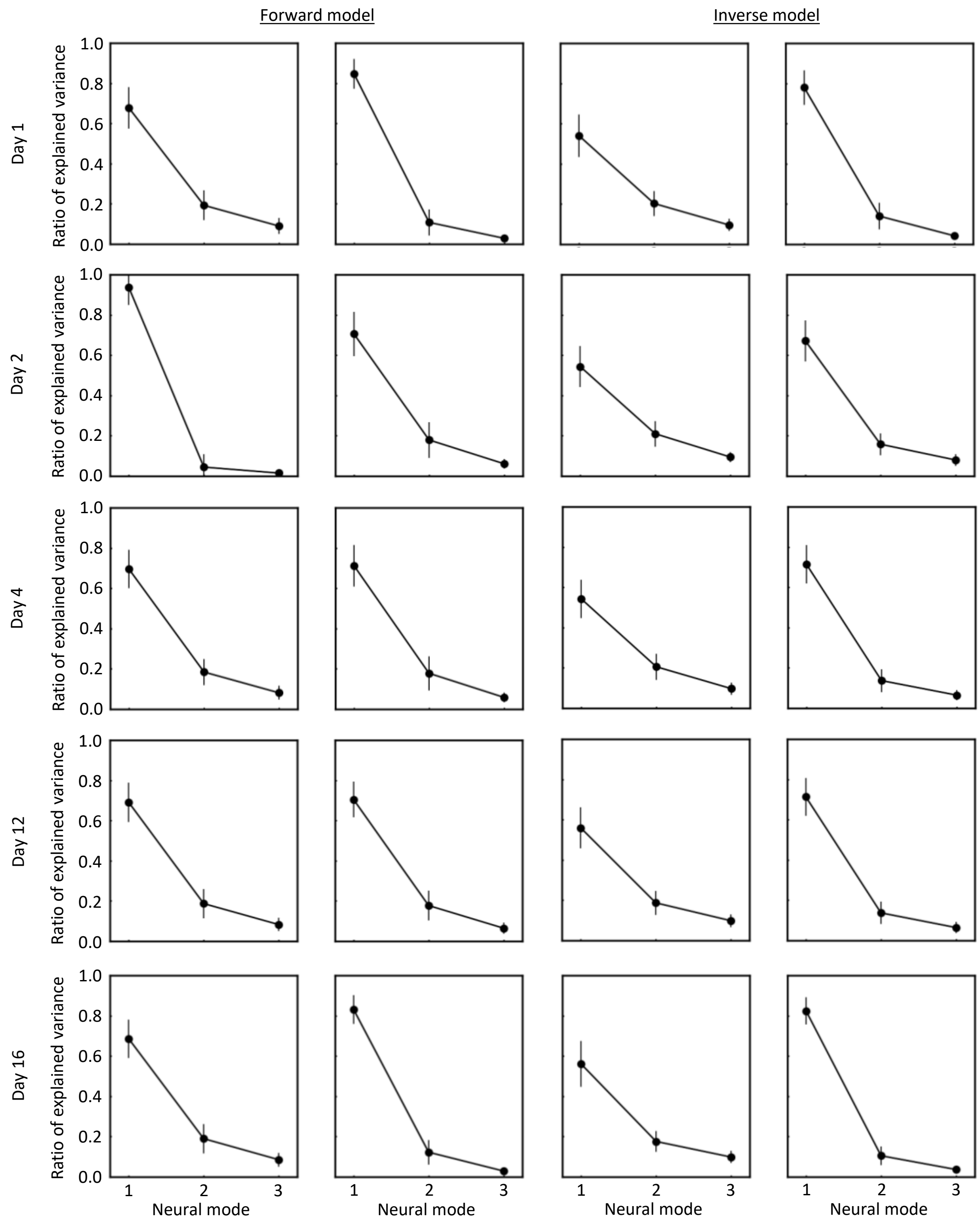


**Supplementary Fig. 5: Identification of neural modes using principal component analysis.** Scree plot of the percentage of variance explained by each principal component (neural model) demonstrates that three components capture most of the variance in the original screw ECoG (1st column), reconstructed screw ECoG (2nd column), original LFP (3rd column), and reconstructed LFP (4th column) across days (error bars, s.d.; Day 1, n = 68; Day 2, n = 49; Day 4, n = 128; Day 12, n = 78; Day 16, n = 135).

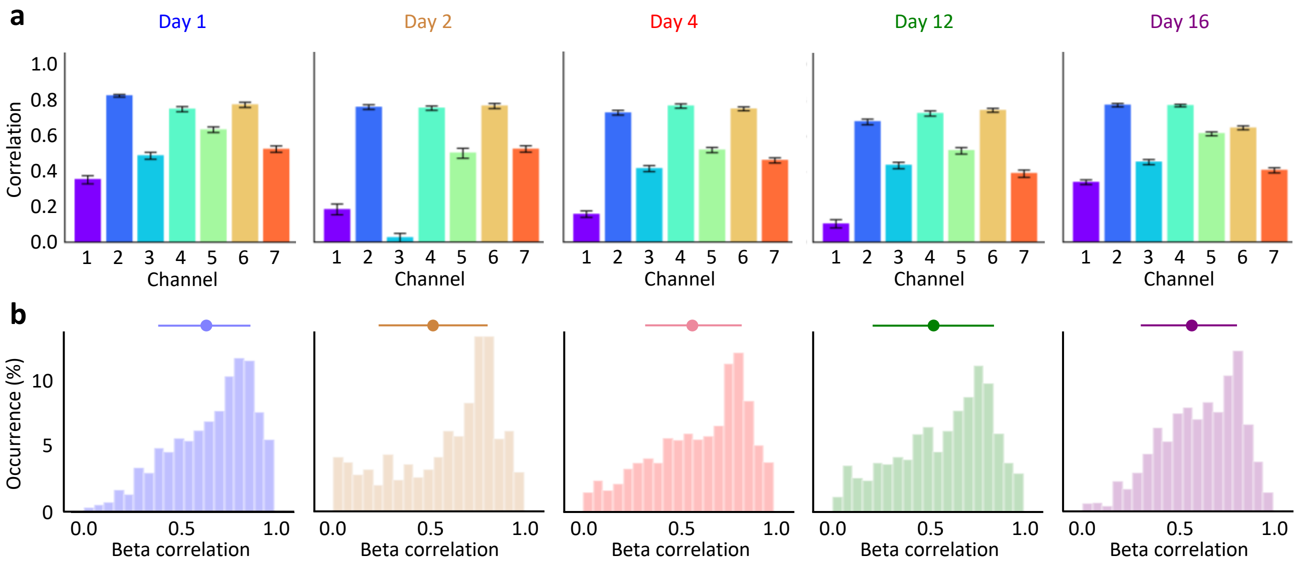


**Supplementary Fig. 6: Stability of forward-NBGNet's performance over time is evaluated with cross-correlation.** (**a**) Average correlation coefficient across the trials and across the days (error bars, s.e.m.; Day 1, n = 150; Day 2, n = 82; Day 4, n = 256; Day 12, n = 165; Day 16, n = 222). Screw ECoG channels correspond to the electrode's layout in **Fig. 2a**. They were almost as correlated across different days as in Day 1. Channel 3 in Day 2 is an outlier where the recordings were unstable. (**b**) Beta correlation between forward-NBGNet's inference and the ground truth screw ECoG across days. Histograms indicates that model predictions are strongly correlated to the ground truths.

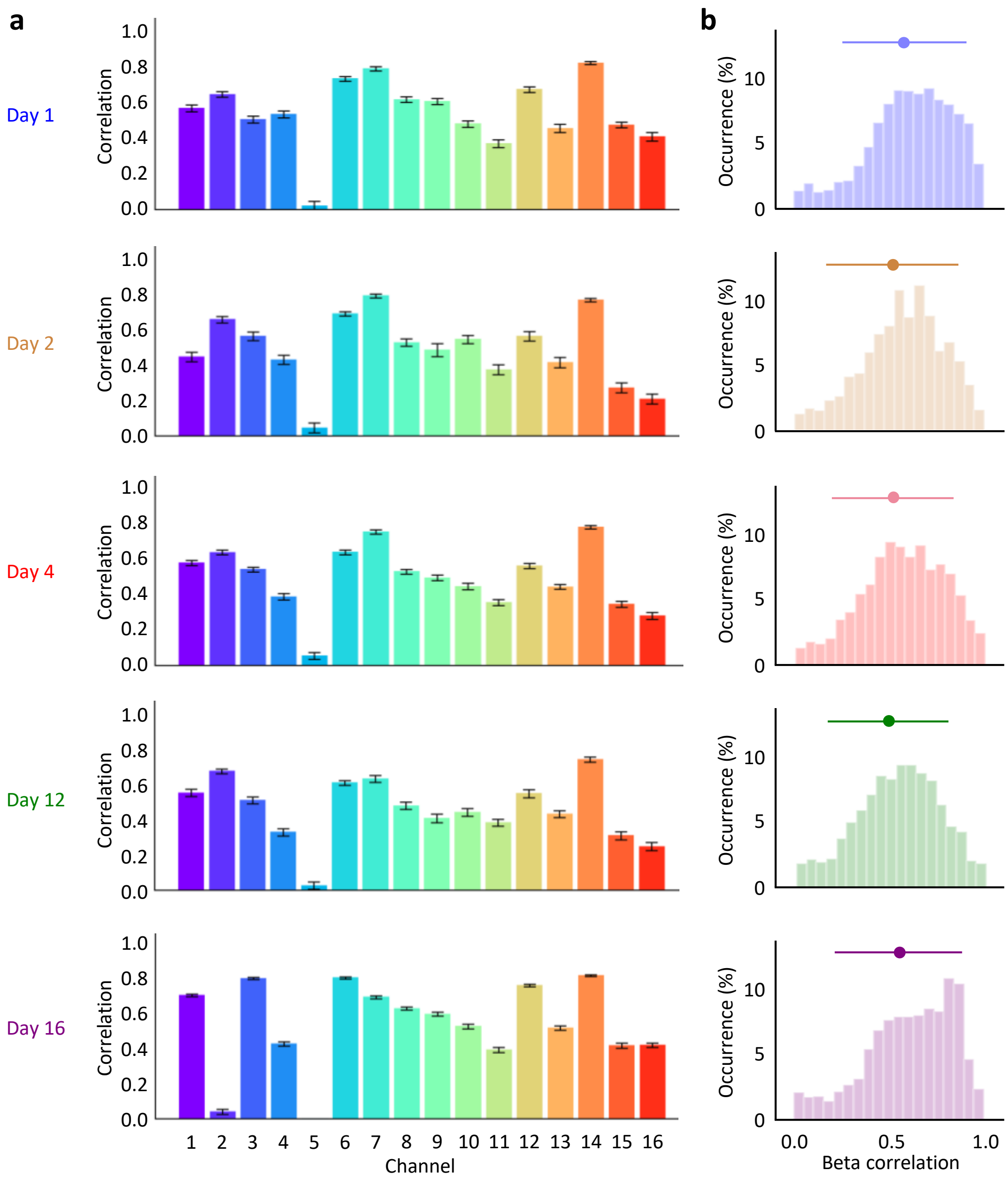


**Supplementary Fig. 7: Stability of inverse-NBGNet's performance over time is evaluated with cross-correlation.** (**a**) Average correlation coefficient across the trials and across the days (error bars, s.e.m.; Day 1, n = 150; Day 2, n = 82; Day 4, n = 256; Day 12, n = 165; Day 16, n = 222). LFP channels correspond to the electrode's layout in **Fig. 2e**. They were almost as correlated across different days as in Day 1. Channel 2 in Day 16 is an outlier where the recordings were unstable. (**b**) Beta correlation between inverse-NBGNet's inference and the ground truth LFP across days. Histograms indicates that model predictions are strongly correlated to the ground truths.

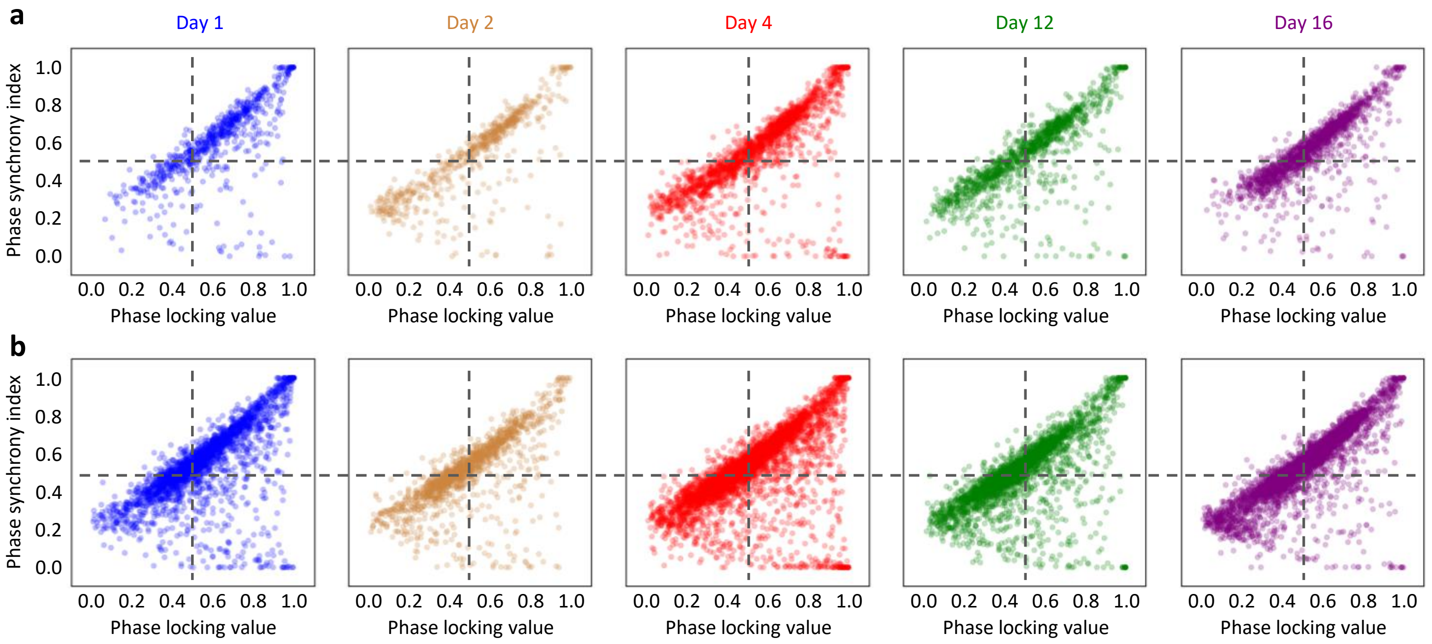


**Supplementary Fig. 8: Stability of NBGNet's performance over time is evaluated with phase synchrony.** (**a**) Phase synchrony index and phase locking value for each channel and trial across the days for forward model (Day 1, n = 1050; Day 2, n = 574; Day 4, n = 1792; Day 12, n = 1155; Day 16, n = 1554). The forward-NBGNet inference and ground-truth screw ECoG were almost as synchronous across different days as in Day 1. (**b**) Phase synchrony index and phase locking value for each channel and trial across the days for inverse model (Day 1, n = 2400; Day 2, n = 1312; Day 4, n = 4096; Day 12, n = 2640; Day 16, n = 3552). The inverse-NBGNet inference and ground-truth LFP were almost as synchronous across different days as in Day 1.

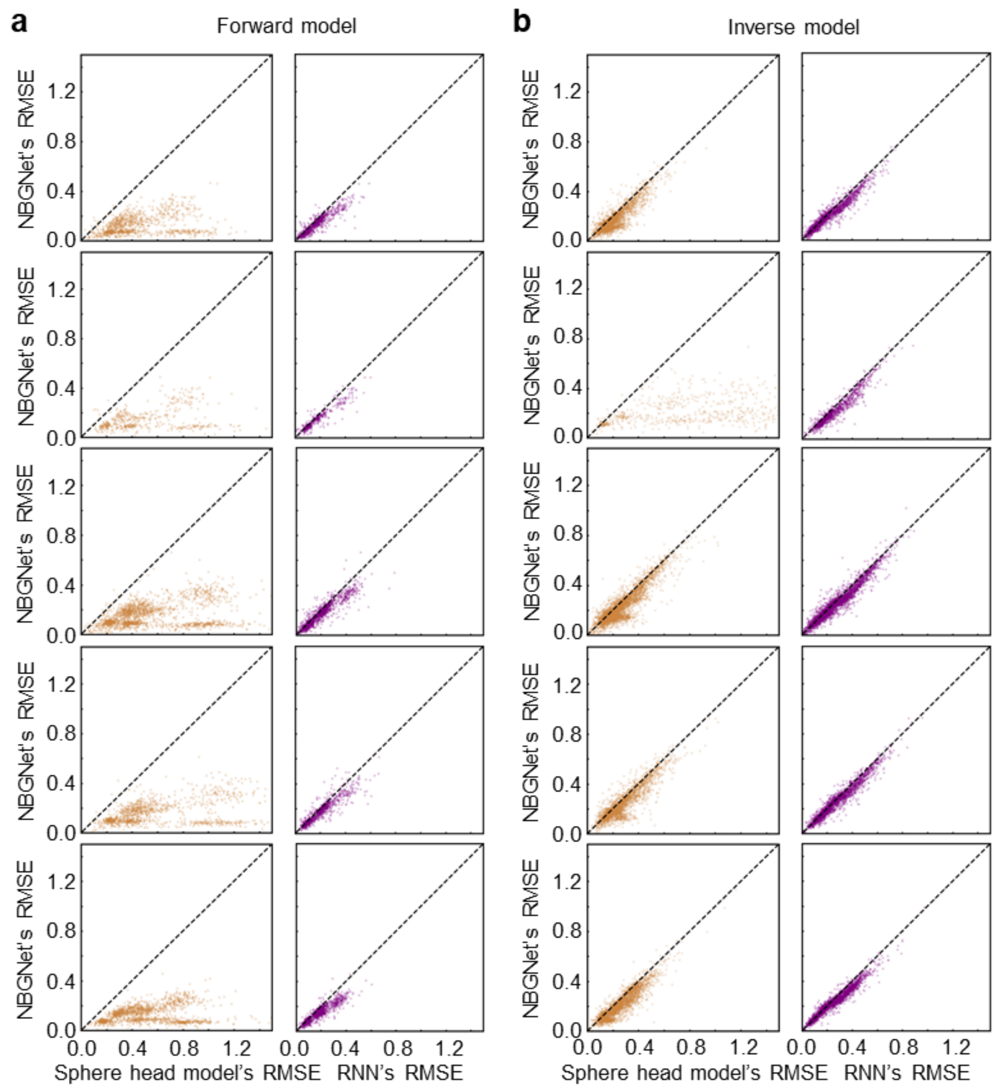


**Supplementary Fig. 9: The performance of NBGNet, sphere head model and RNN was evaluated by RMSE.** (**a**) Two dimensional (NBGNet's RMSE versus sphere head model's RMSE or RNN's RMSE) scattered plots for forward model. The scattered points were below the black dashed bisection line indicating that NBGNet outperformed two other methods with lower RMSE. (**b**) Same as **a** for inverse model. The scattered points were below the black dashed bisection line indicating that NBGNet outperformed two other methods with lower RMSE.

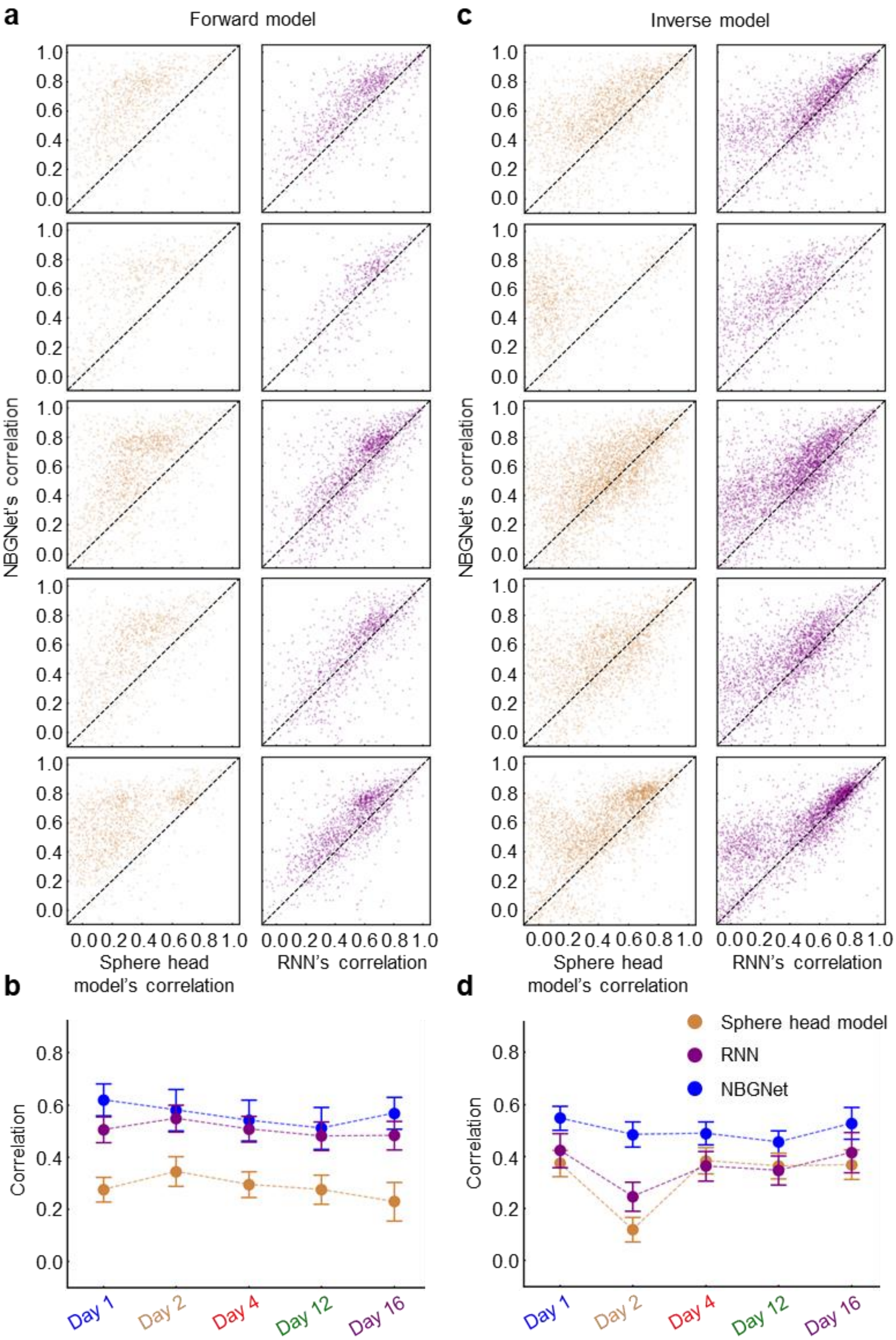


**Supplementary Fig. 10: The performance of NBGNet, sphere head model and RNN was evaluated by cross-correlation.** (**a**) Two dimensional (NBGNet's correlation versus sphere head model's correlation or RNN's correlation) scattered plots for forward model. The scattered points were above the black dashed bisection line indicating that NBGNet outperformed two other methods with higher correlation. (**b**) Scatter plot of average correlation obtained from different methods across days showing no difference in the comparison between each other for forward model (error bars, s.e.m.; n = 7). (**c**) Same as **a** for inverse model. The scattered points were above the black dashed bisection line indicating that NBGNet outperformed two other methods with higher correlation. (**d**) Same as **b** for inverse model (error bars, s.e.m.; n = 16).

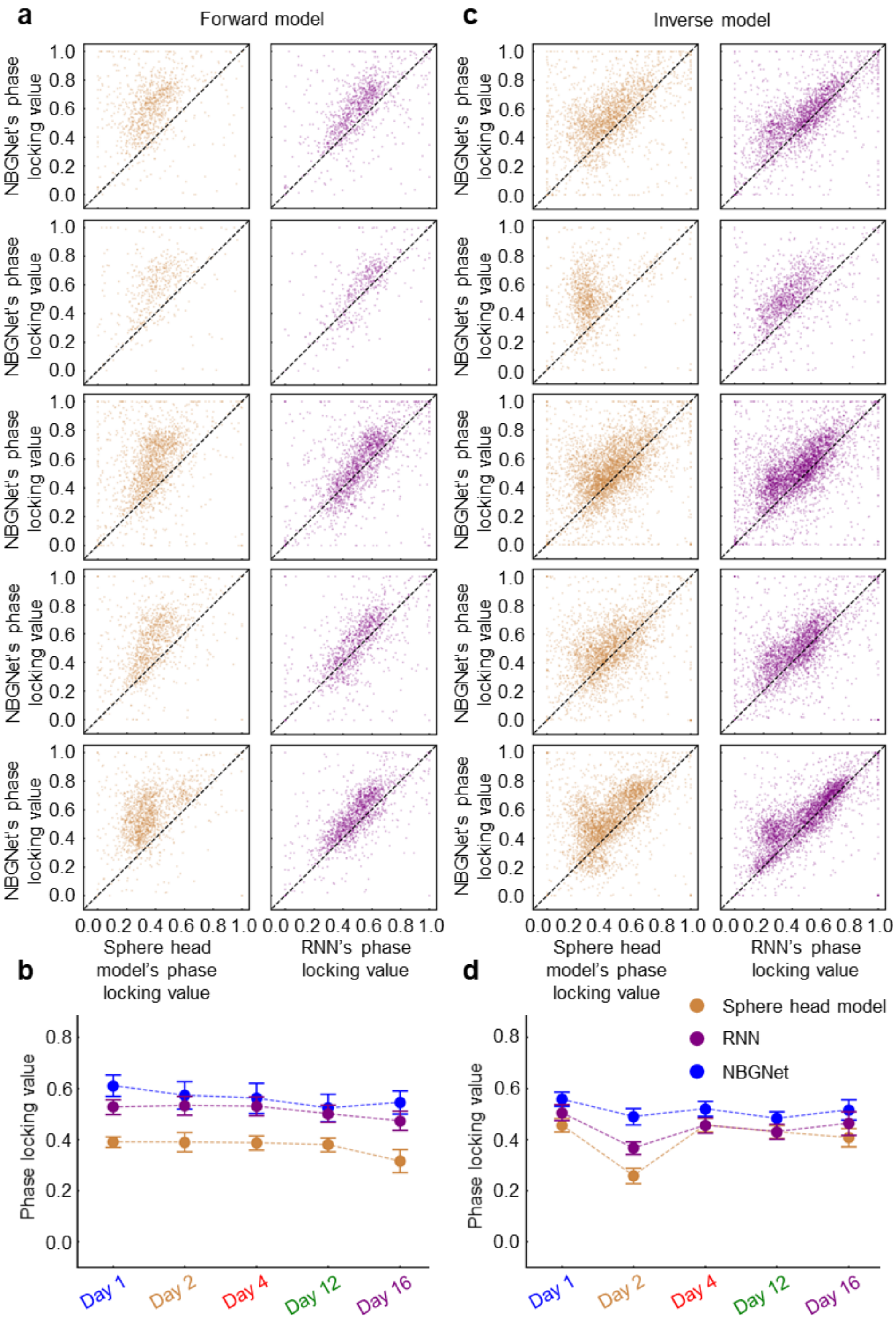


**Supplementary Fig. 11: The performance of NBGNet, sphere head model and RNN was evaluated by phase locking value.** (**a**) Two dimensional (NBGNet's phase locking value versus sphere head model's phase locking value or RNN's phase locking value) scattered plots for forward model. The scattered points were above the black dashed bisection line indicating that NBGNet outperformed two other methods with stronger phase similarity. (**b**) Scatter plot of average phase locking value obtained from different methods across days showing no difference in the comparison between each other for forward model (error bars, s.e.m.; n = 7). (**c**) Same as **a** for inverse model. The scattered points were above the black dashed bisection line indicating that NBGNet outperformed two other methods with stronger phase similarity. (**d**) Same as **b** for inverse model (error bars, s.e.m.; n = 16).

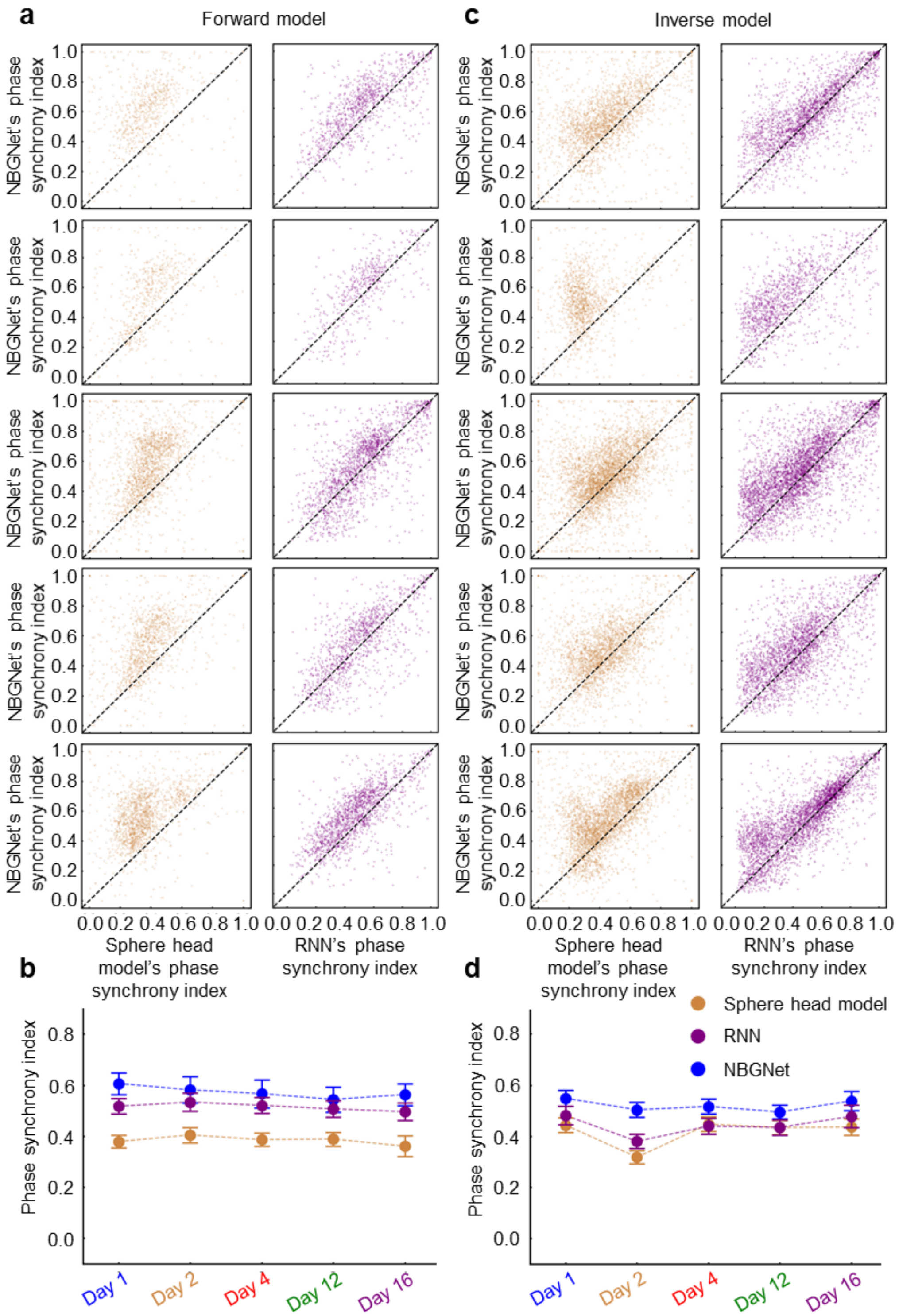


**Supplementary Fig. 12: The performance of NBGNet, sphere head model and RNN was evaluated by phase synchrony index.** (**a**) Two dimensional (NBGNet's phase synchrony index versus sphere head model's phase synchrony index or RNN's phase synchrony index) scattered plots for forward model. The scattered points were above the black dashed bisection line indicating that NBGNet outperformed two other methods with stronger phase synchrony. (**b**) Scatter plot of average phase synchrony index obtained from different methods across days showing no difference in the comparison between each other for forward model (error bars, s.e.m.; n = 7). (**c**) Same as **a** for inverse model. The scattered points were above the black dashed bisection line indicating that NBGNet outperformed two other methods with stronger phase synchrony. (**d**) Same as **b** for inverse model (error bars, s.e.m.; n = 16).

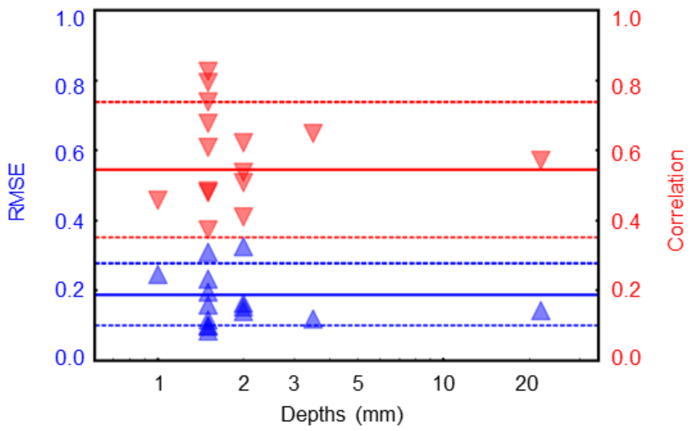


**Supplementary Fig. 13: NBGNet's performance is depth-independent.** Two dimensional (RMSE and cross-correlation versus depths of the electrodes) scattered plots for inverse model indicate that the deepest electrode (> 20 mm) has similar performance with other channels. Solid line, mean without considering the deepest channel. Dashed lines, +/- one s.d. without considering the deepest channel.

**Supplementary Table 1: Screw ECoG electrode layout.**

| **Channel number** | 1 | 2 | 3 | 4 | 5 | 6 | 7 |
|---|---|---|---|---|---|---|---|
| **Hemisphere** | Right | Left | | | | | |
| **Brain region** | S1 | PMv | S1 | lPFC | M1 | mPFC | S1 |

**Supplementary Table 2: LFP electrode layout.**

| **Channel number** | 1 | 2 | 3 | 4 | 5 | 6 | 7 | 8 |
|---|---|---|---|---|---|---|---|---|
| **Hemisphere** | Left | | | | | | | |
| **Brain region** | vmPFC | preSMA | FEF | SMA | PMd | PMd | M1 | PMv |
| **Depth (mm)** | 22.0 | 3.5 | 2.0 | 2.0 | 1.5 | 1.5 | 1.5 | 2.0 |
| **Channel number** | 9 | 10 | 11 | 12 | 13 | 14 | 15 | 16 |
| **Hemisphere** | Left | | | | | | | |
| **Brain region** | M1 | M1 | M1 | M1 | M1 | SMA | M1 | S1 |
| **Depth (mm)** | 1.5 | 1.5 | 1.5 | 1.5 | 1.0 | 1.5 | 1.5 | 2.0 |

**Supplementary Table 3: Selected features for movement behavior decoding (* 5 features selected from each group; ** 10 features selected when considering both groups simultaneously)**

Please see the separate table file.

**Supplementary Table 4: Parameters for sphere head model.**

| **Label** | **Tissue** | **Radius (mm)** | **σ (S/m)** |
|---|---|---|---|
| 1 | Brain | 27.88 | 0.33 |
| 2 | Cerebrospinal fluid | 28.24 | 1.65 |
| 3 | Skull | 30.00 | 0.00825 |
| 4 | Scalp | 31.76 | 0.33 |

### Supplementary Discussion

**Model performance on LFP channel 5.** We observed a consistently poor performance on LFP channel 5, PMd, across days for inverse-NBGNet in **Fig. 2e, f**, **Fig. 6d,** and **Supplementary Fig. 7**. Due to noise on this channel, it provided less stable recordings as compared with other channels. It is thus unsurprisingly that inverse-NBGNet does not infer well the neural activity recorded at LFP channel 5 since there is no underlying stable dynamics to uncover. Although this channel does seem to be exceptional in terms of the poor performance, visual inspection of the raw data and other preliminary analyses (e.g., power spectral density) did not provide enough definitive evidence for it to be excluded *a priori* from the core analyses and thus it is included in all the analyses presented in this manuscript.
